# Kinase inhibitor-induced cell-type specific vacuole formation in the absence of canonical ATG5-dependent autophagy

**DOI:** 10.1101/2023.08.27.554994

**Authors:** Susan Jose, Himanshi Sharma, Janki Insan, Khushboo Sharma, Varun Arora, Sonam Dhamija, Nabil Eid, Manoj B. Menon

## Abstract

Pyridinyl imidazole class p38 MAPKα/β (MAPK14/MAPK11) inhibitors including SB202190 have been shown to induce a cell-type specific defective autophagy response resulting in micron-scale vacuole formation, autophagy-dependent death, and tumor growth suppression *in vivo.* We had earlier shown that this is an off-target effect of SB202190. Here we provide evidence that the cell-type specific vacuole formation is independent of canonical autophagy pathway. While SB202190 seems to interfere with autophagic flux in many cell lines in parallel to vacuolation, autophagy-deficient DU-145 cells and CRISPR/Cas9 gene-edited ATG5 knockout A549 cells also undergo vacuolation upon SB202190 treatment. Late-endosomal GTPase RAB7 colocalizes with these compartments and RAB7 GTP-binding seems to be essential for SB202190-induced vacuolation. RAB7 is a driver of tumor progression and interfering with RAB7-positive endo/lysosomal compartments may enhance cytotoxicity. A screen for modulators of SB202190-induced vacuolation revealed molecules including multi-kinase inhibitor Sorafenib as inhibitor of vacuolation and sorafenib co-treatment enhanced the cytotoxicity of SB202190. Moreover VE-821, an ATR kinase inhibitor was found to phenocopy the cell-type specific vacuolation response of SB202190. To identify the factors determining the cell-type specificity of the vacuolation response induced by SB-compounds and VE-821, we compared the transcriptomics data from vacuole forming and non-vacuole forming cancer cell lines and identified a gene expression signature which may define sensitivity of cancer cells to these small-molecule kinase inhibitors. Further analyses using the small molecule tools and the gene signature discovered here, could reveal novel mechanisms regulating this interesting phenotype relevant to anti-cancer therapy.

## Introduction

Small molecule kinase inhibitors are sought after tools for therapeutic intervention against diverse pathologies including cancer, inflammation and metabolic disorders (reviewed in (1)) and have been developed and tested against cancer and inflammation (2, 3). The discovery of p38 mitogen-activated protein kinases (MAPKs) have been closely linked with the development of kinase inhibitors to target chronic inflammatory disorders. Pyridinyl imidazole class p38 MAPKα/β (MAPK14/MAPK11) inhibitors including SB202190 and SB203580 have facilitated extensive research on the physiological roles of p38 signaling. Despite widespread use of SB202190/SB203580 as specific MAPK14/MAPK11 inhibitors, there are several additional kinase targets known for these molecules including BRAF, CK1, GAK, GSK3β, LCK, PDK1, RAF1, RIPK2 and TGFBR2 (4–6). In addition, these SB compounds have also been shown to target non-kinase proteins including mediators of inflammation. We had earlier shown that SB202190 could effectively and specifically target BRAF-V600E mutant oncogenic kinase to suppress ERK1/2 activity in colorectal cancer and melanoma cells (7). A recent study has verified the use of these pyridinyl imidazole class inhibitors for dual targeting of BRAF and mTORC1 pathways against melanoma (8). Moreover, sensitivity to SB202190 treatment was shown to be effective in predicting BRAF-activating mutations in patient-derived colorectal cancer organoids (9).

Autophagy is a catabolic process wherein cytoplasmic contents including organelles are sequestrated into autophagosomes, followed by subsequent fusion with lysosomes and degradation of contents in the autolysosomal compartments (10, 11). High rates of basal autophagy are prevalent in several cancers, which provide them resistance to metabolic stress, making autophagy a target for anti-cancer therapy (12). Nutrient deprivation is the strongest activator of autophagy and different nutrient sensing pathways tightly regulate autophagy and cell death under nutrient deprived conditions. While mTOR signaling is the primary regulator of autophagy initiation in mammalian cells, various stages of autophagy are regulated by p38, ERK1/2, AMPK and the PI3K-III (phosphatidylinositol 3-kinase, class III) complex (13–15). The PI3K-III complex consisting of BECN1 protein regulates both initiation and maturation of autophagosomes, integrates signals from multiple signaling pathways and also regulates the balance between cell death and survival (16). Initiation of autophagy in response to nutrient or energy deprivation requires assembly and activation of the ULK1 (unc-51-like kinase 1) complex, which leads to the formation of a phagophore, facilitated by the ATG5-ATG12-ATG16 complex mediated lipidation of LC3/ATG8 protein. The phagophore expand and engulf autophagy cargo and form a double membrane autophagosomes which finally undergo maturation and fusion with lysosomes to form autolysosomes, degrading the contents and releasing vital metabolites (17). Canonical autophagy is abrogated in ATG5- and ATG7-deficient cells, however recent evidence suggests the presence of a non-canonical pathway where RAB9 GTPase mediates the elongation of the initial autophagosomal membrane, independent of ATG5-mediated LC3-lipid conjugation (18).

Mammalian cells undergo cytoplasmic vacuolation in response to diverse stimuli and this has often been associated with cell death (19). While autophagy may be involved in vacuole formation, several pathogens as well as phytochemicals induce autophagy-independent vacuoles. Paraptosis, a caspase-independent cell death pathway with accumulation of cytoplasmic vacuoles originating from endoplasmic reticulum and swollen mitochondria has been targeted for cancer therapy (20, 21). Methuosis is another vacuolation dependent mode of programmed cell death where vacuoles originate from macropinosomes and swollen endosomes (22, 23). SB202190 was originally shown to induce vacuoles specifically in colorectal cancer cells (24, 25). The micron-scale vacuoles induced by SB202190, and related pyridinyl imidazole compounds were characterised as autolysosomes, originating from a defective autophagy response (7, 25). While most of the vacuolation-dependent programmed cell death pathways can be targeted for anti-tumor strategies, the cell-type specificity of SB202190-induced vacuolation makes it more interesting.

Autophagy often plays a complex role in cancer. This can be attributed to opposing roles for pathway components in diverse aspects of tumour initiation, progression and therapy resistance. Autophagy inhibitors have entered preclinical and clinical evaluation against several forms of cancers (26). Studies using SB202190 had previously established a pro-tumorigenic role for p38α/MAPK14 in colon cancer and SB202190 treatment was shown to induce cell-type specific autophagy-dependent cell death (ACD)(24, 25). This initiation of ACD was attributed to a transcriptional shift from hypoxia-dependent to Foxo-3A-dependent pro-autophagic gene expression and it switches to apoptosis upon inhibition of autophagy (9). In our previous work we could show that the cell-type specific effects of SB202190 are independent of p38 inhibition and transcriptional reprogramming (7). The SB202190-induced vacuoles are unusually large to be autophagosomes and are reminiscent of a defect in autophagy clearance, resulting in enlarged vacuoles. We further investigated this cell-type specific phenotype and present evidence here that the vacuolation induced by SB compounds do not require canonical autophagy. Moreover, we identified small molecule modulators of SB202190-induced vacuolation. While multi-kinase inhibitor Sorafenib was among the molecules which inhibited SB202190-induced vacuolation, VE-821, an ATR inhibitor was found to phenocopy the cell-type specific vacuole forming effects of SB202190. We also discovered a gene expression signature which defines susceptibility of cancer cell lines to SB202190-induced autophagy-independent vacuolation. These findings may reveal novel mechanisms and nodes for targeting autophagy-dependent cell death beyond the ambit of canonical autophagy pathways.

## Materials and Methods

### Reagents

SB202190 (Cayman,#1000399), SB220025 (Sigma, S9070), Bafilomycin A1 (Cayman, #11038), BIRB-796 (Cayman, #10460), Leupeptin (Cayman, #14026), Chloroquine (Sigma, C6628), Apilimod (MedChemExpress/MCE, HY-14644), Vacuolin-1 (MCE, HY-118630), Ceralasertib (MCE, HY-19323), Elimusertib (MCE, HY-101566), CID-1067700 (MCE, HY-101566) and all other inhibitor stocks were prepared in DMSO. Antibodies for LC3A/B (#12741), ATG5 (#12994), ATG12 (#4180), Phospho-Chk1 (Ser345)(#2348) were from Cell Signalling Technology, MA, USA. Rab7 (#A12308) and Rab11a (#A3251) antibodies were from Abclonal, MA, USA, RAB9 antibody from Invitrogen (#MA3-067) and p62/SQSTM1 (#BB-AB0130) was from Bio-Bharati Life Science, West Bengal, India and GAPDH (#GTX100118) was from GeneTex, CA, USA. Goat Anti-Rabbit IgG(H+L) (HRP conjugated, #111-035-045) was from Jackson ImmunoResearch Laboratories, PA, USA and Goat Anti-Rabbit IgG (H+L)(AF488 conjugated, E-AB-1055) was from Elabscience, Wuhan, China. Phalloidin iFluor™ 647 Conjugate (#20555) was from Cayman chemicals. DMEM (#11965092), FBS (#10270106), Trypsin-EDTA (#15400054), Antibiotic-antimycotic solution (#15240062), Goat anti-Rabbit IgG (H+L)-Alexa Fluor™ 488 (#A11070) and Goat anti-Mouse IgG (H+L)-Alexa Fluor™ 568 (#A11019) were purchased from Thermo Fisher Scientific, MA, USA. DMEM/F12 (#AL215A), EBSS (#TL1110), Sodium pyruvate solution (#TCL015), Bovine Serum Albumin (BSA) (#MB083) and Dimethyl Sulfoxide (DMSO) (#TC185) were obtained from HiMedia Laboratories, Mumbai, India. Details of the inhibitors used in the screen is provided in Supplementary table S1. Other biochemicals were obtained from Sisco Research Laboratories (SRL), Mumbai, India.

### Cell culture and Treatments

HCT116, HT29, 293T, L929, HeLa, U-87MG and MDA-MB231 cells were maintained in DMEM with 10% FCS. Media for HCT116, 4T1, C2C12, Huh7, NIH-3T3, LN229, HeLa, DU145 and MCF7 were additionally supplemented with 1mM Sodium pyruvate solution. NCI-H460 was cultured in RPMI with 10% FCS. MCF10A was maintained in DMEM/F12 + 5% Horse serum + 20 ng/ml EGF + 0.5mg/ml hydrocortisone + 100 ng/ml cholera toxin + 10 µg/ml insulin and A549, PC-3 and HaCaT cells were maintained in DMEM/F12 with 10% FCS. All cell lines other than MCF10A were maintained in the presence of 1x antibiotic-antimycotic solution (#15240062, Thermo). All Cell lines were maintained at 37 °C with 5% CO2. Cells were treated with inhibitors at indicated concentrations in 12 well plates. DMSO was used as solvent control. All treatments were performed in normal growth medium for the respective cell lines. SB202190 was used at 10 µM and BafA1 at 100 nM concentration, unless stated otherwise in the figure legends. For monitoring ATR activity, cells were stimulated with UV-C (10J/m^2^) and were recovered for 1 h. A549 cells were transfected with GFP-RAB7^wt^, GFP-RAB7^Q67L^ and GFP-RAB7^T22N^ expression vectors (kind gift from Dr. Amit Tuli, CSIR-IMTECH, Chandigarh) (27) using Lipofectamine-3000 reagent following manufacturer’s protocol. Cells were transfected for 6 h in antibiotic free medium, followed by medium change and two hours recovery before SB202190-treatment for 18 h followed by microscopy with a standard fluorescent microscope.

To delete *ATG5* gene from A549 cells, all in one CRISPR/Cas9 vector was used as described previously (28). Previously described sgRNA sequences for *ATG5* deletion (29, 30) were synthesised as DNA oligos and cloned in pLKO5.U6-chiRNA.EF1a.hSpCas9-P2A-PAC.pre vector (kind gift from Dr. Dirk Heckl, MHH, Hannover) and lentiviral vectors were packaged using HEK293T cells, as described previously (31). After transduction, cells were selected with puromycin (2ug/ml) and single cell clones were screened for ATG5 expression by Western blotting. Control and KO single-cell clones were isolated and further characterized.

### Immunostaining and Microscopy

For experiments, cells were seeded in 12 well plates and treated with small molecule inhibitors. Live phase-contrast images were captured using camera attached to DIM150 microscope (Debro or Quasmo). Inhibitor treatments were for 24 h if not mentioned otherwise. Percentage vacuolation was calculated manually (Figure 3D) or by Image-J based image analyses as described below. For Immuno-fluorescence staining, cells were seeded on poly-L-lysine precoated glass cover slips. After treatment, cells were washed with 1X-PBS and fixed with 4% paraformaldehyde (PFA) for 2 mins at room-temperature (RT), followed by 20mins on ice. Cells were permeabilized with 0.25% Triton X-100 for 20 min at RT and blocked using 3% BSA-PBS for 1h at 4°C. Primary antibodies were used at a 1:100 to 1:200 dilution in 1% BSA–PBS for 2 h at RT. Alexa-488 and Alexa-568 labelled secondary antibodies were used at 1:250-1:1000 dilution in 1% BSA-PBS at RT for 45 min, followed by DAPI (500ng/ml in PBS) staining for 10 min at RT. Phalloidin iFluor™ 647 (1:200) was included in the secondary antibody mixture wherever indicated. Imaging was performed using a Leica TCS SP8 confocal microscope with standard settings.

### Image analyses for quantification of vacuolation

To quantify vacuolation in A549 cells in response to SB202190 and other inhibitors (Figure 2G & 4A), ImageJ v1.54d (https://imagej.nih.gov/ij/) was used. For each image, the contrast was enhanced (saturated pixels=0.35%) and the green channel 8-bit image was extracted using the RGB-split tool. The threshold for each image was set manually (between 195-255 raw intensity values) to select vacuoles and the number of vacuoles was counted using the *Analyze Particle* feature (size=100-250, circularity=0.80-1.00). All these steps were recorded as a macro and run to analyse multiple images from each inhibitor treatment (4 images per treatment). To normalize the findings across experiments, the data was presented as percentage vacuolation, with the number of vacuoles in the SB202190-treated wells considered as 100%.

### Western Immunoblotting

Cells were lysed directly in SDS gel loading dye after treatments and lysates separated on 12-14% SDS-PAGE gels, transferred to nitrocellulose membranes, blocked with 5% non-fat dry milk for 1h and probed with primary antibodies for 16h and secondary antibodies for 1h. Blots were developed using ECL reagents (Westarnova 2.0, Cyanagen) and bands were visualized using ImageQuant LAS 500 (Cytiva) or Fusion Solo 6S chemiluminescence imager (Vilber).

### Differential expression analyses and visualization

RNAseq read counts for cell lines in CCLE (Cancer Cell Line Encyclopedia) available on the Cancer Dependency Portal (DepMap, Release 23Q2) were downloaded on 24.07.2023 (32, 33). From the dataset, 15 cell lines with experimentally verified data for vacuolation phenotype in response to SB202190 were selected for further analyses. The RSEM values were adjusted to the nearest integer and only the genes with non-zero expression counts in ≥5 out of the 15 cell lines were used for differential gene expression analysis using DESeq2 (v1.38.3) in R (v4.2.3) (34, 35). DESeq results were sorted in the order of adjusted p-values and VST-normalized values for the top fifty genes were used to plot a heatmap in R. Volcano plot depicting the differentially expressed genes (|log2FC|>2 and *p-*adj <0.01) and the boxplots of VST normalized values, representing expression of selected genes were plotted using ggplot2 (v3.4.3)(36). The overlapping off-targets of SB202190 and VE-821 were obtained by comparing previously published datasets (37, 38). In case of SB202190, kinomescan targets with dissociation constants (Kd) <5 μM were considered and for VE-821, all tested human kinases are shown (Supplementary Figure 9A).

### Cell viability assays

Cell viability was measured in triplicates in 96-well plates using MTT [3-(4,5-dimethylthiazol-2-yl)-2,5-diphenyltetrazolium bromide]. Cells were seeded at 4000 cells/well density in 96 well plates and were treated for 72hrs with indicated inhibitors. After the treatments, cells were incubated with 10ul MTT solution (5mg/ml in 1x Phosphate-buffered saline, #58945, SRL) for additional 2-3 h at 37°C. Formazan crystals were dissolved in 100ul DMSO and absorbance taken at 570nm, with 690nm as reference using a plate reader (Cytation 5, BioTek).

### Gene expression analyses by real-time qPCR

RNA was isolated using the NucleoSpin RNA extraction kit (Macherey & Nagel) and reverse transcribed (Bio Bharati, BB-E0043) using random hexamer primers according to the manufacturer’s instructions. qRT-PCRs were run on a CFX96 (Bio-Rad) device using TB Green Premix Ex Taq II (Tli RNase H Plus, Takara) and the gene expression was normalized to Cyclophilin and presented. Sequences of the primers used are listed in Supplementary Table S2.

### Statistics and Reproducibility

All immunoblot and microscopy results are representative of at least three independent results. Quantitative assays are performed in triplicates and data presented as mean values ± SD using Microsoft Excel. Students 2-tailed t-tests (unpaired, unequal variance) were used to check the statistical significance of differences between sample groups in the quantitative assays. For differential gene expression data, statistical analyses and calculation of adjusted p-values were performed using default options in the DESeq2 (v1.38.3) package. Additional details are mentioned in figure legends. The statistics and source data including actual *p*-values are presented in Supplementary Table S4.

## Results

### SB202190-induces cell-type specific vacuoles which accompanies altered autophagy flux and is inhibited by bafilomycin A1

We have previously shown that SB202190 and related pyridinyl imidazole family kinase inhibitors of p38α (MAPK14)/p38β (MAPK11) induce cell-type specific vacuolation attributed to defective autophagosomes (7, 39). To understand the mechanisms leading to this strong phenotype, we expanded our studies to include more cell lines, in addition to those reported earlier (Table 1). While 22 out of 33 cell lines tested till date showed vacuolation upon treatment with SB202190 (10 μM for 24 h), different variants of HeLa cells gave different results. The variations in HeLa is consistent with a recent study which compared 14 strains of HeLa cells and identified high phenotypic variability, attributed to proteotypic variability (40). Previously, we have shown that the vacuole formation is due to off-target effects of the pyridinyl imidazole family of p38 MAPK inhibitors including SB202190 and SB203580 and is not phenocopied by more specific p38 inhibitors like BIRB-796 or VX-745 (7). Interestingly, SB220025, a closely related pyridinyl imidazole-based p38 inhibitor of the same family also did not induce vacuoles (Figure 1A & 1B). This may hint towards structural differences in the pharmacophores determining the binding of the SB-family compounds to the autophagy/vacuole formation-relevant targets (Figure 1B).

**Figure 1.**
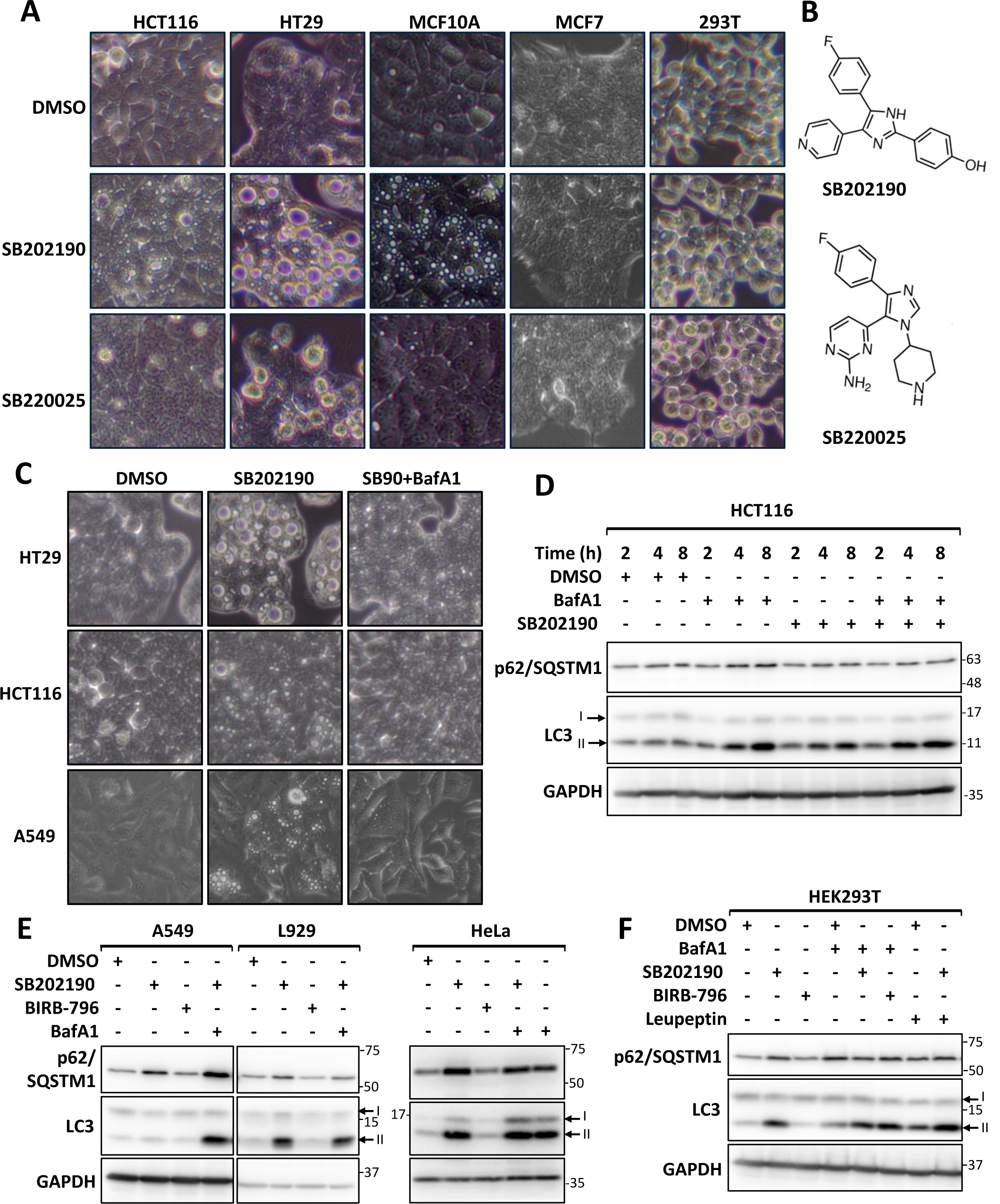
SB202190-induced cell type specific vacuolation accompanies suppression of autophagy flux and is inhibited by Bafilomycin A1. **A.** Indicated cell lines were treated with SB202190 (10 µM), SB220025 (10 µM) or solvent control (DMSO) for 24 h and images were acquired using a 20x phase-contrast objective. **B.** Structures of pyridinyl imidazole class inhibitors SB202190 (upper panel) and SB220025 (lower panel). **C.** Effect of Bafilomycin A1 (BafA1) treatment on SB202190-induced vacuolation in HT29, HCT116 and A549 cells. **D.** HCT116 cells were treated with BafA1 and/or SB202190 for indicated time-points and lysates were probed with p62 and LC3 antibodies to monitor autophagy flux. GAPDH is shown as control. **E.** A549, HeLa and L929 cells were treated with indicated small molecules for 24 h and lysates were probed for LC3, p62 and GAPDH as in panel D. **F.** HEK293T cells were treated as indicated for 24 h and lysates were analysed by immunoblotting.

**Table 1.** Cell-type specificity of SB202190-induced vacuole formation.

| **No** | **Cell line** | **Species** | **Cell type** | **Vacuoles** |
| --- | --- | --- | --- | --- |
| 1 | **A549** | Human | lung carcinoma | + |
| 2 | AGS | Human | gastric adenocarcinoma | + |
| 3 | BHK21 | Hamster | adult kidney fibroblast | + |
| 4 | **C2C12** | Mouse | myoblast | - |
| 5 | Caco-2 Bbe | Human | colorectal adenocarcinoma | + |
| 6 | **DU-145** | Human | prostate carcinoma | + |
| 7 | **HaCaT** | Human | keratinocyte | - |
| 8 | **HCT 116** | Human | colorectal adenocarcinoma | + |
| 9 | **HEK293T** | Human | embryonic kidney | - |
| 10 | **HeLa** | Human | cervical adenocarcinoma | -/+ |
| 11 | hMSC | Human | primary mesenchymal stem cells | + |
| 12 | **HT29** | Human | colorectal adenocarcinoma | + |
| 13 | **Huh7** | Human | hepatocellular carcinoma | - |
| 14 | HUVEC | Human | primary endothelial cells | + |
| 15 | IEC6 | Rat | small intestinal epithelium | + |
| 16 | **L929** | Mouse | fibrosarcoma | + |
| 17 | **LN229** | Human | glioblastoma | + |
| 18 | **MCF-10A** | Human | Mammary epithelial | + |
| 19 | **MCF7** | Human | breast adenocarcinoma | - |
| 20 | **MDA-MB231** | Human | breast carcinoma | + |
| 21 | MEF-T | Mouse | embryonic fibroblast | + |
| 22 | **NCI-H460** | Human | lung carcinoma | + |
| 23 | **NIH 3T3** | Mouse | embryonic fibroblast | - |
| 24 | NMuMG | Mouse | mammary epithelial | + |
| 25 | **PC-3** | Human | prostate adenocarcinoma | - |
| 26 | RAW 264.7 | Mouse | monocytic | + |
| 27 | RGM1 | Rat | gastric epithelium | + |
| 28 | Sh-SY5Y | Human | neuroblastoma | - |
| 29 | SW480 | Human | colorectal adenocarcinoma | + |
| 30 | **U87MG** | Human | glioblastoma | + |
| 31 | WM1617 | Human | melanoma | - |
| 32 | WM793 | Human | melanoma | - |
| 33 | **4T1** | Mouse | breast carcinoma | + |

Notes: The cell lines tested in the present study are shown in bold. Others have been reported in previous studies (7, 39). -/+ indicate presence of different results from different sublines of the same cell line.

We then investigated the effect of the lysosomal acidification inhibitor Bafilomycin-A1 (BafA1). SB202190-induced vacuole formation was inhibited by BafA1 treatment (Figure 1C). In addition, we monitored the levels of the autophagy substrate p62/SQSTM1 and the autophagosome marker LC3-II (lipid conjugated LC3) by immunoblotting. SB202190 treatment led to significant accumulation of the autophagy markers in A549, L929 and HEK293T cells, which was further enhanced by BafA1 treatment (Figure 1D-F). The effect of BafA1 and SB202190 on p62/LC3-II accumulation showed differences in kinetics and strength between the different cell lines tested (Figure 1D-F & Supplementary Figure S1A). Consistent with the previous results, the pan-p38 inhibitor BIRB-796 did not alter the autophagy flux (Figure 1E & 1F). Autophagy flux analyses using the protease inhibitor leupeptin also led to an increase in LC3-II and p62 levels similar to BafA1 treatment (Figure 1F). Interestingly, the effect of SB202190 on the autophagy flux was not connected to vacuole formation, since it was also observed in non-vacuole forming HEK293T cells (Figure 1F). This indicates the possibility that vacuole formation could be independent of alterations in the autophagy flux.

### SB202190 induces vacuoles in inherently autophagy-deficient DU-145 prostate cancer cells

In our effort to expand the cell type specific aspect of the vacuolation phenotype, we also tested the effect of SB202190 on the prostate cancer cell lines DU-145 and PC3. While PC3 cells did not display significant vacuole formation upon SB202190-treatment (Table 1), DU-145 cells formed vacuoles which were inhibited upon BafA1 co-treatment (Figure 2A). Consistent with the other cell lines, SB220025 did not induce vacuoles in this cell line. Interestingly, due to the presence of a mutation, which leads to the expression of a non-functional splice variant of ATG5, DU-145 cells have previously been characterised as an autophagy-deficient cell line (41). We verified this by immunoblotting and there were no ATG5-ATG12 and LC3-II conjugates detectable in this cell line indicating the absence of active autophagosome biogenesis (Figure 2B). This unexpected discovery argues against the previous conclusions that the SB202190-induced micron-scale vacuoles are of autophagosome origin.

**Figure 2.**
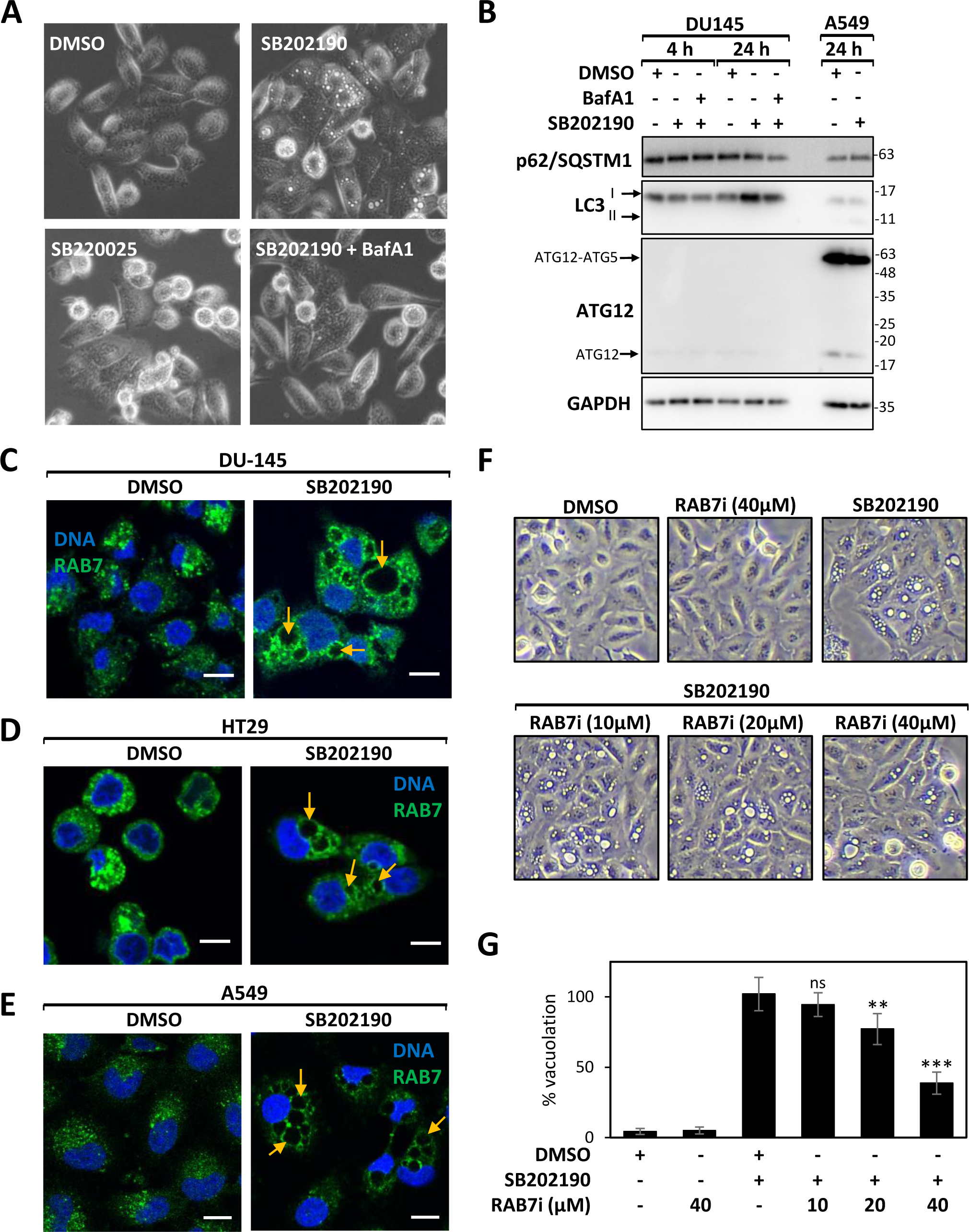
Autophagy-deficient DU-145 cells induce RAB7-positive vacuoles in response to SB202190. **A.** DU-145 cells were treated as indicated for 24 h and imaged using a 20x phase contrast objective to observe vacuoles. **B.** Lysates from DU-145 cells were probed with p62, LC3 and ATG12 antibodies to monitor autophagy. ATG12 antibody predominantly detects ATG12-ATG5 conjugates (55 kDa) in autophagy proficient A549 cells. GAPDH is shown as control. **C - E.** Confocal microscopy analyses of RAB7 localisation in DU-145 (C), HT29 (D) and A549 (E) cells treated with SB202190 or solvent control (DMSO) for 24 h. **F & G.** A549 cells were treated with SB202190 (10 µM) in the presence or absence of indicated concentrations of RAB7 inhibitor (RAB7i) for 24 h. Representative images (F) and percentage vacuolation quantified (G)( ** denotes p<0.01; ***denotes p<0.001 and ns denotes p>0.05) are shown.

To further understand the identity of these SB202190-induced acidic vacuoles, we performed immunofluorescence analyses of RAB7 and RAB11 in SB202190-treated cells. RAB7 is a small GTPase associated with late endosomes, involved in autophagosome and late-endosome fusion with lysosomes (42, 43). The large vacuoles induced by SB202190 were clearly labelled by RAB7 in both autophagy-proficient HT29/A549 cells and autophagy-deficient DU-145 cells (Figure 2C-2E). Similar immunofluorescence analyses for RAB11, a marker for recycling endosomes (44) did not reveal any redistribution upon SB202190-treatment (Supplementary Figure S1B). We also tested the colocalization of the RAB7-positive vacuoles with RAB9, a marker of non-canonical autophagosomes. Interestingly, SB202190 treatment in A549 cells led to enhanced appearance of RAB9 containing puncta, but these were distinct from the micron-scale RAB7+ vesicles (Supplementary Figure S1C). To verify whether RAB7 is actively involved in the formation of these vesicles, we treated cells with CID-1067700 (RAB7i), a GTPase inhibitor which strongly inhibits RAB7 nucleotide binding (45, 46). Pretreatment with RAB7i dose dependently inhibited SB202190-induced vacuolation (Figure 2F-2H). Moreover, a constitutively GTP-bound mutant of GFP-RAB7 (Q67L) showed colocalization with larger micron-scale vacuoles, compared to the wild-type protein (Supplementary Figure S2A) and vacuolation was suppressed in cells transfected with dominant negative non-GTP binding mutant RAB7 (GFP-RAB7^T22N^) (Supplementary Figure S2B). Thus, SB202190 induced vacuoles are swollen RAB7 positive vesicular compartments, which are sensitive to BafA1, dependent on RAB7 GTP binding, but not obligately dependent on autophagy.

### Gene-edited A549 cells confirm autophagy-independent formation of SB202190-induced vacuoles

To conclusively prove the lack of involvement of canonical autophagy pathways in the generation of SB202190-induced micron-scale vacuoles, we performed CRISPR/Cas9-mediated gene editing in vacuole forming A549 lung adenocarcinoma cells. CRISPR/Cas9 all-in-one vector expressing Cas9 and sgRNA against *ATG5* gene was used to generate canonical autophagy-deficient cells (Supplementary Figure S3). After gene deletion, two independent clonal cell lines of each genotype (wild type – WT, ATG5-knockout – KO) were compared. The *ATG5* KO cells displayed higher basal levels of LC3-I and p62/SQSTM1 (Figure 3A & 3B). Moreover, upon induction of autophagy by amino-acid starvation, there was no detectable change in autophagic flux in the *ATG5*-KO clones compared to the control cell lines (Figure 3A). When SB202190 was used to alter the autophagy pathway, we observed similar results, with the *ATG5*-KO cell lines neither displaying p62/SQSTM1 degradation, nor inducing LC3-II conjugation (Figure 3B). We then monitored SB202190-induced vacuole formation in these cell lines. While there were clonal differences in the extent of vacuole formation, both WT and *ATG5*-KO A549 cell lines displayed significant vacuolation in response to 24 h of SB202190 treatment (Figure 3C & 3D). Immunofluorescence analyses again revealed RAB7 labelled compartments in A549 cells upon SB202190-treatment independent of the presence of ATG5 (Figure 3E). These data conclusively prove that even in an autophagy proficient cell line like A549, SB202190-induced vacuole formation is independent of the canonical autophagy pathway. Unlike colon cancer cells (7, 24), we did not observe autophagy-related gene expression changes in A549 cells (Supplementary Figure S4), and, therefore the role of the *ATG5*-deletion on gene expression could not be analysed further.

**Figure 3.**
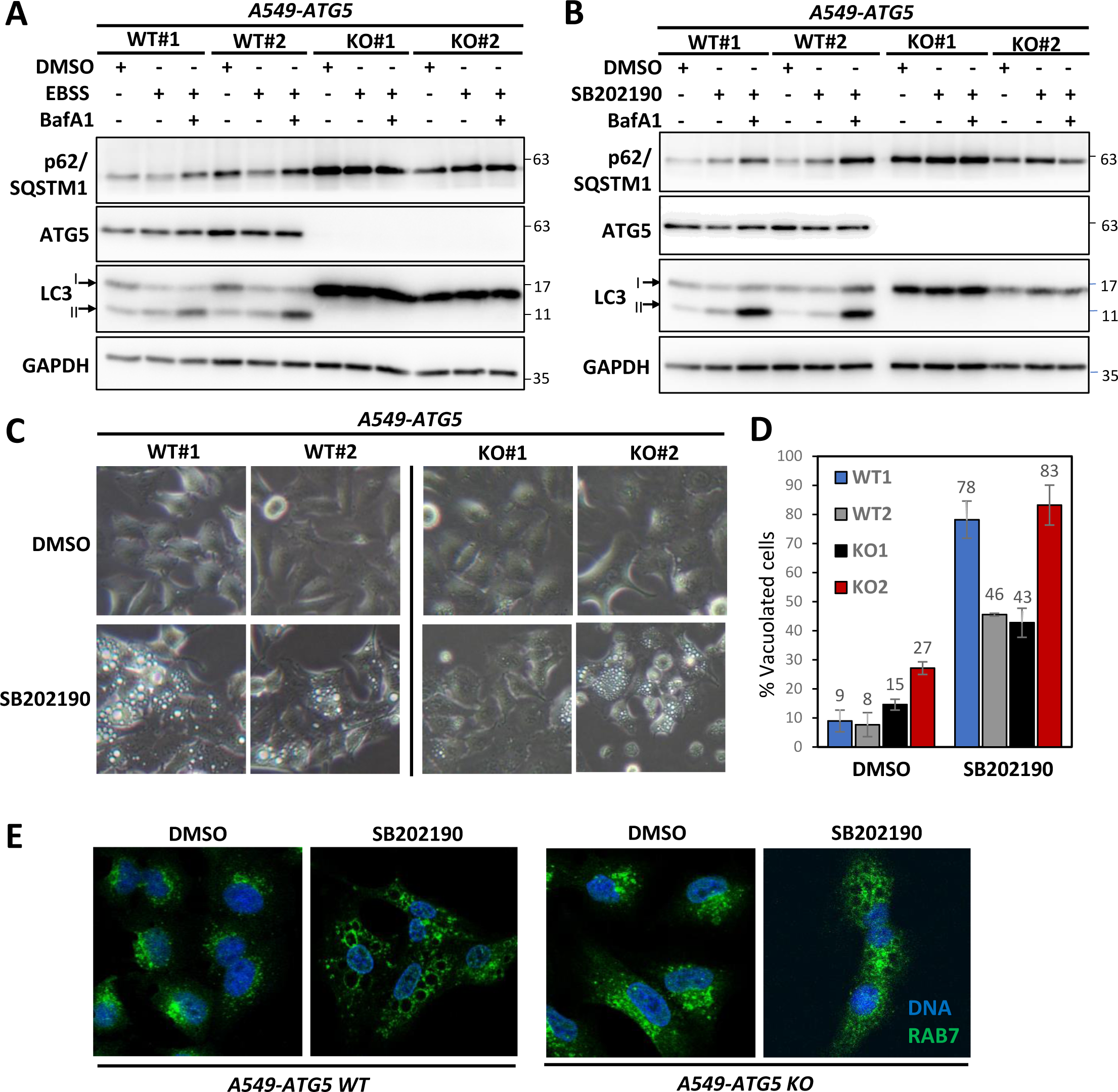
Canonical autophagy and LC3-conjugation is dispensable for SB202190-induced vacuolation. **A.** EBSS (Earle’s Balanced Salt Solution) treatment-induced autophagy was monitored in *ATG5 deficient*-A549 cells by probing the degradation of autophagy adapter and substrate p62 and the autophagy marker – lipid conjugated lower band of LC3 (LC3-II). Bafilomycin A1 is used as a control to determine autophagic flux and DMSO as solvent control. EBSS was for 3 hrs, while control cells were in complete DMEM/F12 medium. ATG5 antibody mainly detects ATG12-ATG5 conjugates in cells visible at ∼55 kDa. **B.** Western Blot analysis shows LC3-II accumulation in WT cells co-treated with SB202190 (20 µM) and BafA1. **C & D.** Clone-specific differences are seen in the number of vacuoles but vacuoles were observed in both WT and ATG5 KO cells(C). **E.** SB202190-induced vacuoles are GTPase Rab7 positive and are independent of the presence or absence of ATG5 in the cells.

### SB202190-induced vacuole formation is modulated by multiple small molecule inhibitors including the multi-kinase inhibitor Sorafenib

Since the vacuoles are not of autophagy origin, we investigated other possible pathways involved in the biogenesis of SB202190-induced vacuoles. For this, we co-treated A549 cells with a panel of diverse small molecule inhibitors including modulators of cytoskeleton/vesicular trafficking and kinase inhibitors (Supplementary Table S1). As expected, lysosomotropic agents like BafA1 and chloroquine inhibited the vacuole formation induced by SB202190 (Figure 4A). Interestingly, among the 54 small molecules tested, we also obtained significant suppression of vacuole formation by several additional compounds including kinase inhibitors (Figure 4A & Supplementary Figure S5). While suppression of vacuolation was associated with cell death and major morphological changes in case of some molecules (indicated by asterisks in Figure 4A), the compounds MS-049 (PRMT4/6 inhibitor), Gefitinib (EGFR tyrosine kinase inhibitor) and Sorafenib (multiple tyrosine kinase inhibitor) inhibited vacuole formation in A549 cells without apparent toxicity upon co-treatment with SB202190 for 24 h (Figure 4B). Interestingly, four compounds including (i) pan-p38 inhibitor BIRB-796 (BIRB796), (ii) the calcineurin phosphatase inhibitor Cyclosporin-A (CsA), (iii) ferroptosis inducer Erastin and (iv) ATR kinase inhibitor VE-821 significantly enhanced SB202190-induced vacuole formation (Figure 4B). We also performed similar experiments in HT29 cells, where the vacuole formation was more pronounced at 24 h (Figure 4C). While Sorafenib inhibited the extent of SB202190-induced vacuolation in HT29, the effects were less pronounced. The inhibitory effect of MS-049 and gefitinib was restricted to a reduction in the large-sized vacuoles (Figure 4C). This confirms the idea that there are cell-type specific differences in the pathways contributing to the vacuolation phenotype. Previous studies have shown that SB202190 induced significant viability loss in vacuole forming cell lines and there was apoptotic death upon suppression of vacuolation/autophagy. Interestingly, treating cells with a combination of sorafenib and SB202190 led to an increase in cell death compared to sorafenib alone (Supplementary Figure S6A-S6C). Consistent with this, an earlier study had demonstrated the additive effect of SB202190 and sorafenib against colon cancer *in vivo* (47). It should be noted that MCF7 and Huh7, two non-vacuole forming cell lines also showed sensitivity to SB202190 treatment (Supplementary Figure S6D). Moreover, all tested cell lines displayed varying levels of sensitivity to other p38 MAPK inhibitors, indicating an additional direct role for p38 inhibition in the anti-cancer effect (Supplementary Figure S6E).

**Figure 4.**
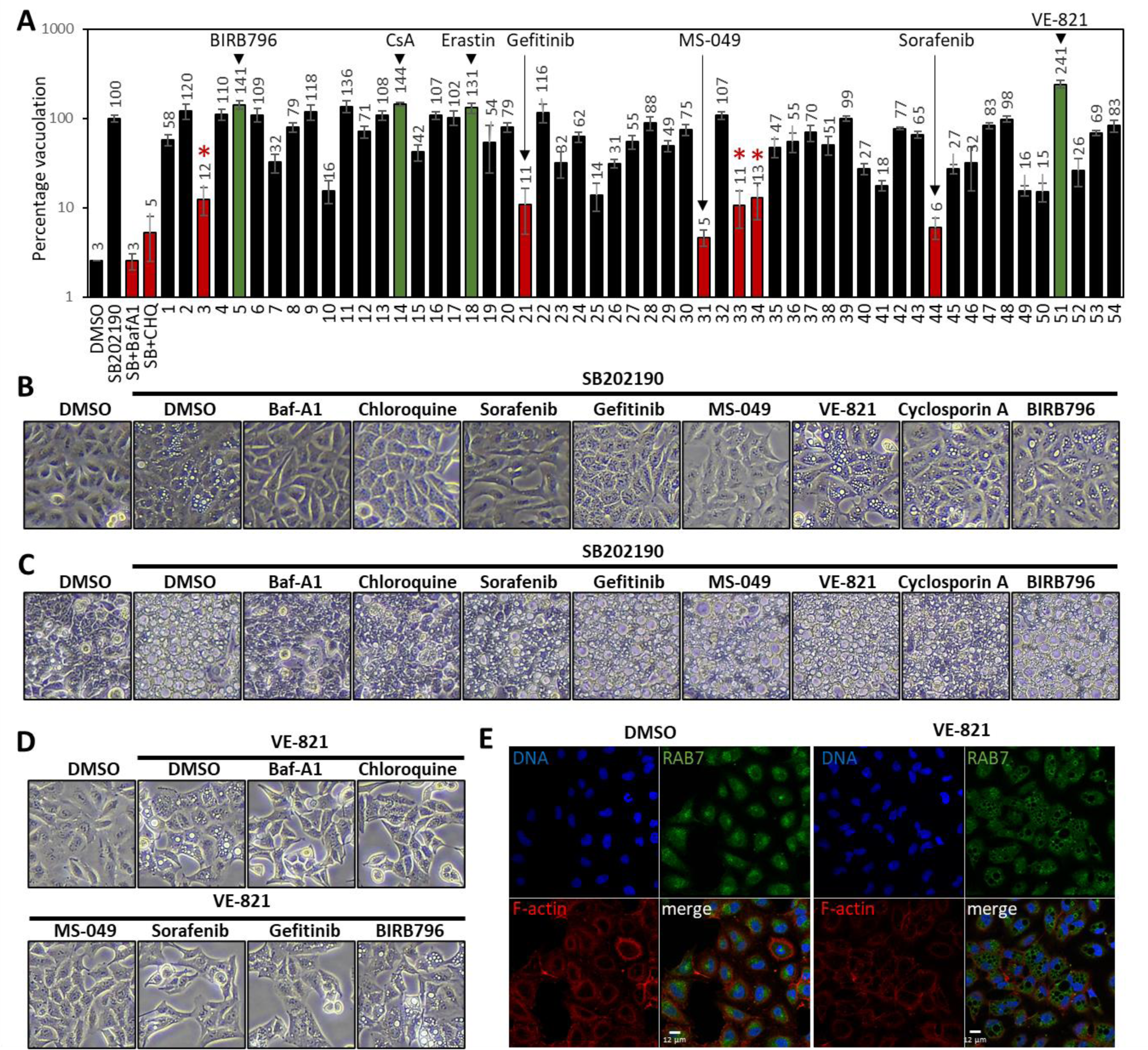
Screen for modulator of SB202190-induced vacuoles identify Sorafenib and VE-821. **A.** To screen for small molecule modulators of SB202190-induced vacuolation, A549 cells were treated with a panel of inhibitors in combination with SB202190 (10 μM) for 24 h. Vacuole formation was quantified and represented taking SB202190 alone treated as 100%. The molecules showing significantly lower or higher vacuolation are indicated with red and green bars respectively. * indicates loss of viability. **B & C.** A549(B) and HT29(C) cells were treated with SB202190 (10 μM) in combination with indicated inhibitors and phase contrast images were taken after 24 h. **D.** A549 cells were treated with VE-821 (10 μM) alone or in combination with indicated inhibitors for 24 h. **E.** A549 cells treated for 24 h with DMSO control or VE-821 (10 μM) were subjected to confocal immunofluorescence analyses to detect RAB7 positive vesicles. DNA (DAPI staining) and F-actin (phalloidin staining) are shown as control.

### VE-821 phenocopies the cell-type specific effects of SB202190 and induce RAB7-positive vacuoles

We then focused on the small molecules which had positively modulated SB202190-induced vacuolation in the screen. When A549 cells were treated with Cyclosporin-A, Erastin or BIRB-796, no micron-scale vacuoles were detected (data not shown). Interestingly, treatment of A549 cells with the ATR inhibitor VE-821 alone was sufficient to induce vacuoles similar to SB202190 (Figure 4D). More importantly, the modulators of SB202190-induced vacuolation identified above had similar effects on VE-821-induced vacuolation, suggesting that these vacuoles are of similar origin. To further support this notion, we monitored RAB7 localisation in VE-821 treated A549 cells. Immunofluorescence staining and confocal microscopy revealed that the VE-821 treated vacuoles were decorated by RAB7 (Figure 4E), similar to the SB202190-induced vacuoles (Figure 2C & 2D). We then investigated whether VE821-induced vacuolation displayed a similar cell-type specificity as SB202190. The cell-type specificity profile of VE-821 perfectly matched with that of SB202190 in all nine cell lines tested (Table 2 and Supplementary Figure S7). These data clearly support the hypothesis that these two inhibitors interfere with the same or similar pathways to induce a cell-type specific enlargement of a RAB7-positive vesicular compartment, independent of canonical ATG5-dependent autophagy. In addition, we observed enhanced SB202190- and VE-821-induced vacuolation when co-treated with the pan p38 inhibitor BIRB-796 indicating a role of p38 inhibition in facilitating vacuolation (Figure 4C & 4D).

**Table 2.** Comparison of cell-type specificity between SB202190 and VE-821-induced vacuole formation.

| **No** | **Cell line** | **SB202190** | **VE-821** |
| --- | --- | --- | --- |
| 1 | A549 | + | + |
| 2 | DU-145 | + | + |
| 3 | HaCaT | - | - |
| 4 | HCT116 | + | + |
| 5 | HEK293T | - | - |
| 6 | HT29 | + | + |
| 7 | MDA-MB231 | + | + |
| 8 | PC-3 | - | - |
| 9 | U87MG | + | + |

### A distinct gene-expression signature defining sensitivity to SB202190-induced vacuole formation

The autophagy interfering activity of VE-821 has been reported previously and is also attributed to off-target effects rather than the inhibition of its primary target ATR (48). To test whether the vacuole-forming effect of VE-821 is also independent of ATR inhibition, we used two other structurally unrelated ATR kinase inhibitors-Ceralasertib and Elimusertib. While both these inhibitors effectively suppressed ATR activity in HT29 and A549 cells as monitored by the inhibition of UV-induced CHEK1/Chk1 phosphorylation, neither of them induced vacuoles in these cells (Supplementary Figure S8). However, it was interesting that SB202190 and VE-821 displayed similar cell type specificity in our studies. SB202190-induced autophagy and colon-cancer specific autophagy-dependent death was originally proposed to be dependent on a gene expression switch between hypoxia-dependent and pro-autophagic Foxo3a-dependent gene expression (24). To understand the gene expression signature which determines the sensitivity of cell lines to formation of these RAB7+ micron-scale vacuoles, we utilized the transcriptomics datasets available from the Cancer Cell Line Encyclopaedia (CCLE) (33). Out of the 23 human cell lines tested for the SB202190-induced vacuolation response, gene expression data was available for 16 cell lines. We excluded HeLa cells from the analyses due to the variations in the vacuolation phenotype associated with different strains of this cell line and the remaining 15 lines included 10 vacuole-forming and 5 non-vacuolating cell lines. We then identified differentially expressed genes between these two groups of cell lines. The analyses revealed an interesting array of genes differentially expressed between the groups of cell lines defined by their susceptibility to SB202190-induced vacuole formation (Figure 5A and 5B). A total of 128 genes (*p value* ≤ 0.01) were differentially expressed between the two groups (Supplementary Table S3), of which 35 were upregulated and 93 downregulated in vacuole-forming cell lines in comparison to the cell lines not susceptible to SB202190-induced vacuolation. Interestingly, the top candidates upregulated in vacuole-negative cell lines included several proteins with vesicular and membrane localisation like LAMP5, AEBP1, RELN etc (Figure 5C). These genes potentially block vacuole formation and define a signature of cancer cell lines based on the basal endo-lysosomal transport and autophagy flux in these cancer cells. In another approach, we identified common off-targets of SB202190 and VE-821 based on previously published inhibitor selectivity datasets (Supplementary Figure S9A). However, these potential common targets of vacuole forming inhibitors were not differentially expressed between vacuolating and non-vacuolating cancer cell lines (Supplementary Figure S9B).

**Figure 5.**
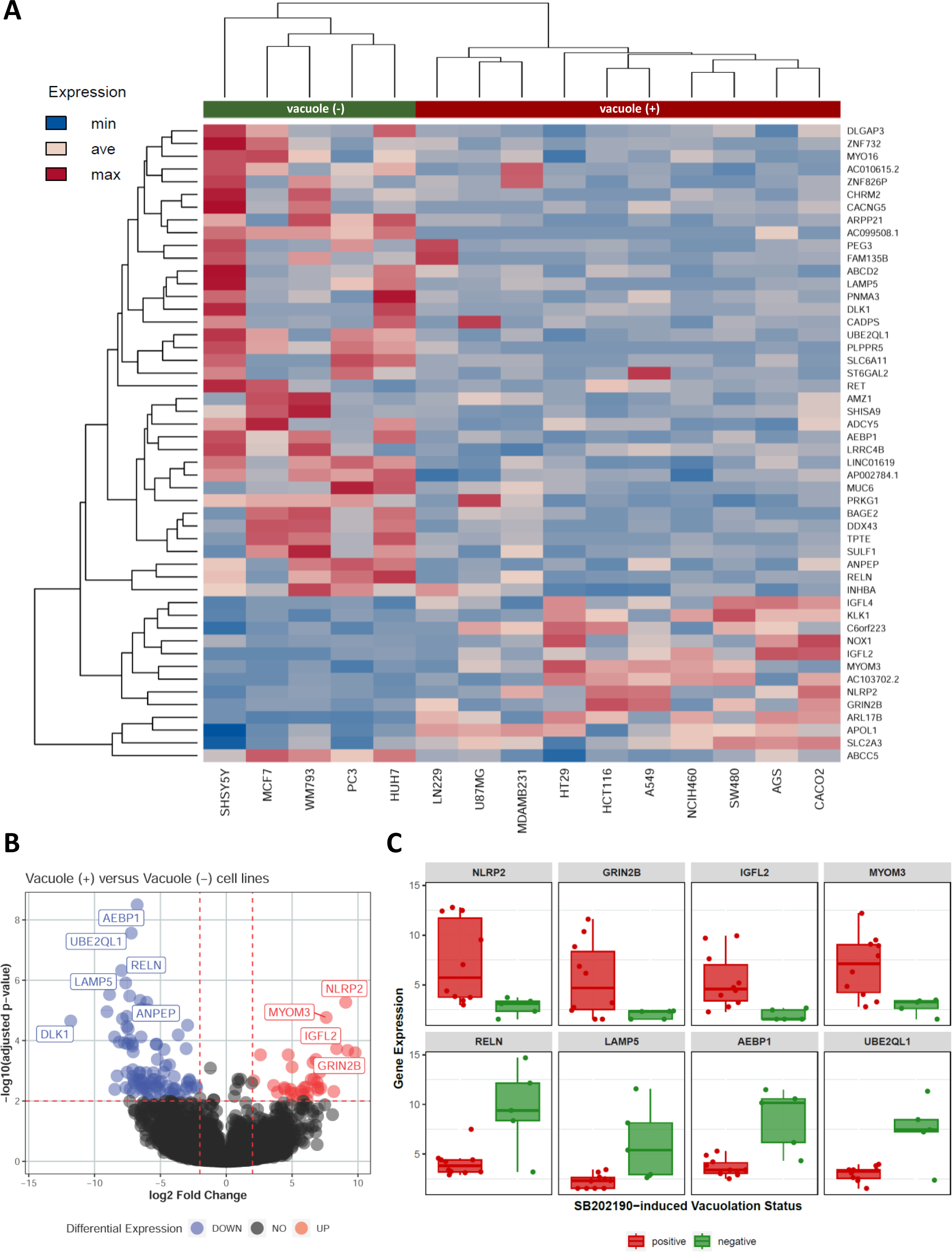
A gene expression signature defining the sensitivity of cancer cell lines to SB202190-induced vacuolation. **A.** Heatmap showing the expression (VST normalized values) of top 50 differentially expressed genes (after sorting by adjusted *p*-values) between the vacuole-forming (vacuole positive) and non-vacuolating (vacuole negative) cell lines. The vacuole-forming cell lines are indicated in red and the non-vacuole forming cell lines are indicated in green. **B.** Volcano plot representing the differential expressed genes identified in the study. The interesting candidate genes showing differential expression are indicated. **C.** Boxplots of VST normalized values, representing expression of select differentially expressed genes across 15 cell lines (red: vacuolation positive and green: vacuolation negative).

## Discussion

SB202190 was originally reported to induce a colon-cancer specific autophagy-dependent cell death, and this was demonstrated to be an effective anti-cancer approach using colon tumor models in mice (24, 25). Reprogramming from a hypoxia driven transcriptional program to Foxo3-dependent gene expression was proposed as the underlying mechanism (24). Our previous work had revealed that SB202190 cell-type specifically influences autophagy flux independent of gene expression changes and inhibition of p38 MAPK activity (7). Here, we have further expanded our analyses and present evidence that the cell type-specific vacuolation response induced by SB202190 proceeds even in the absence of the canonical autophagy mediator ATG5. In the search to identify the signaling pathways facilitating the development of these micron-scale RAB7-positive compartments, we identified small molecules including the multi-kinase inhibitors Sorafenib and Gefitinib and the PRMT4/6 inhibitor MS-049 as inhibitors of SB202190-induced vacuole formation. SB202190 and Sorafenib seems to have an additive effect on inducing cell death in A549 and HT29 cells (Supplementary Figure S6A-6C). Our screen also discovered the ATR kinase inhibitor VE-821 as a small molecule which phenocopies the SB202190-induced cell-type specific vacuolation response. By comparing the transcriptome datasets of vacuole forming and non-vacuole forming cells, we have identified a gene expression signature which may define the sensitivity of human cancer cell lines to SB202190/VE-821 induced vacuolation. These findings (Figure 6) take us closer to the understanding of this strong cell type-specific response affecting vesicular transport and sorting.

**Figure 6.**
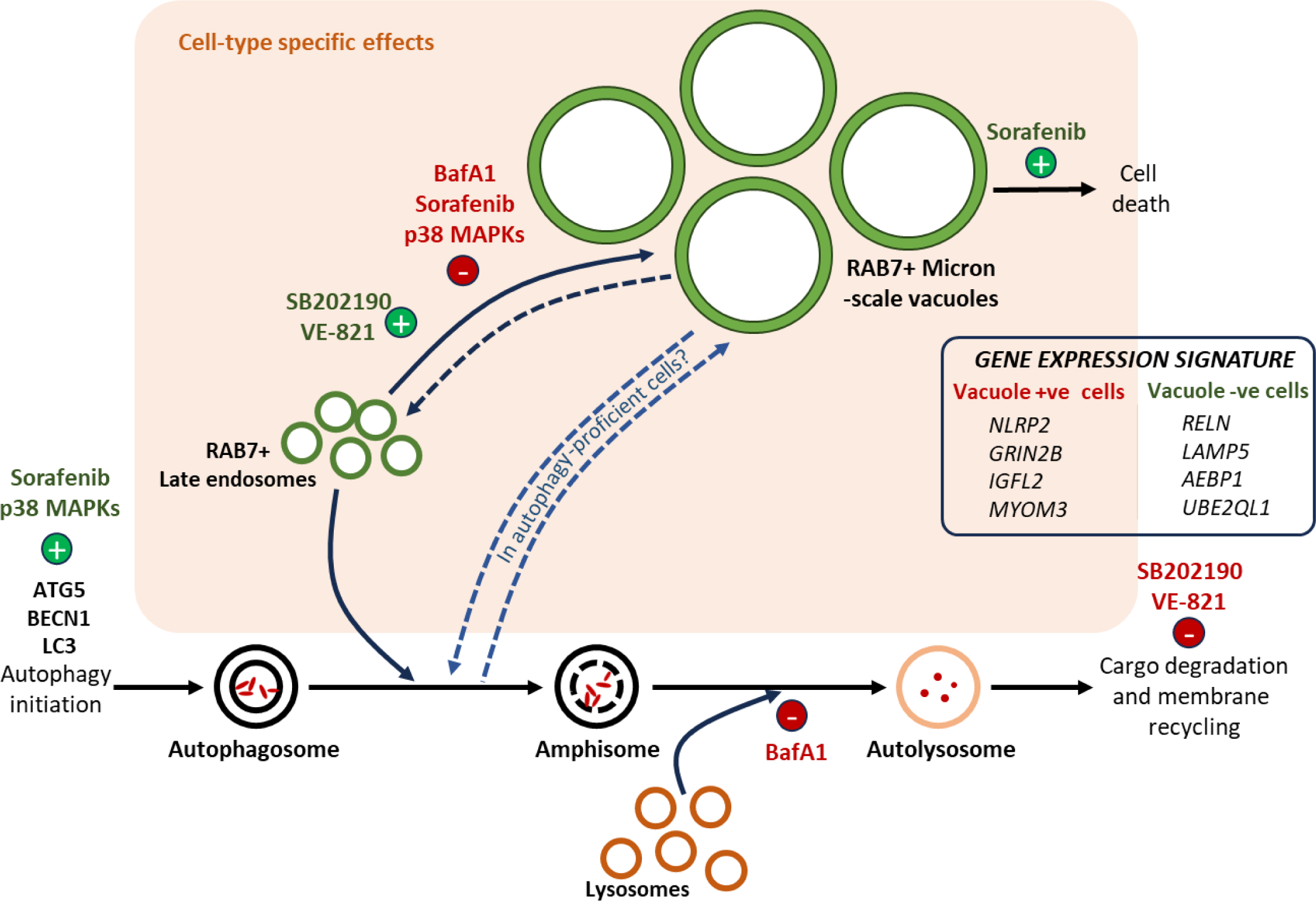
Kinase-inhibitor induced cell-type specific vacuole formation and cytotoxicity. Bottom panel shows normal autophagy pathway wherein autophagosomes arise in response to multiple stressors including nutrient deprivation (ATG5/LC3/BECN1 and the p38 MAPKs are among positive regulators of this process amongst others. Sorafenib has been shown to induce autophagy. Autophagosomes may fuse with late endosomes to give amphisomes and further mature to autolysosomes, after fusion with lysosomes. This results in the degradation of the autophagy cargo and recycling of membrane and nutrients. SB202190/VE-821 kinase inhibitors suppress this autophagic flux. In specific cell-types, partly defined by the gene expression signature (in set), SB202190/VE-821 leads to micron-scale vacuole formation. These vesicles are RAB7-GTP dependent, RAB7+ and may interplay with autophagosomes in autophagy-proficient cells. The process is regulated by additional p38 inhibition and multi-kinase inhibitor sorafenib, in addition to Bafilomycin A1 (BafA1), a molecule known to prevent autolysosome formation/maturation. SB202190/sorafenib co-treatment leads to enhanced cell death in vacuole forming cells.

The pyridinyl imidazole class of MAPK14/p38α-MAPK11/p38β-inhibitors led to the discovery of these kinases as master regulators of inflammation (49). Over the past three decades, the use of SB compounds in cell culture and animal models was a key method in the characterisation of this pathway. Despite studies revealing off-targets (4, 38), these compounds have been essential tools in p38 MAPK research. Considering the autophagy interfering activity of these compounds, we had recommended to avoid the use of these class of p38 inhibitors in characterization of the role of p38 in autophagy research (10, 39). Even though genetic studies have revealed a role for p38 MAPKs and the downstream kinase MK2 in regulation of autophagy (50), several of the phenotypes on autophagy and lysosomal biogenesis reported using SB-compounds can be attributed to these off-target effects (51, 52). Interestingly, SB202190 was shown to induce TFE3/TFEB activation and regulate autophagy and lysosome biogenesis independent of p38 inhibition (53). Recent studies indicate that the effect of SB202190 on TFEB nuclear localisation is mediated by p38-independent EIF2AK3/PERK activation (54). Unlike the micron-scale vacuole formation, these effects on TFEB and PERK activation were not reported to be cell-type specific. We have identified SB220025, an SB-compound closely related to SB202190 as a potential pyridinyl imidazole compound and p38 inhibitor with no effect on the vacuolation response. Interestingly, multiple vacuole-forming and non-vacuolating cell lines showed sensitivity towards a panel of p38 inhibitors in addition to SB202190, reaffirming a direct role for p38 in cancer cell survival (Supplementary Figure S6). In addition, enhancement of vacuolation SB202190 and VE-821-induced vacuolation by BIRB-796 also indicates a crosstalk between p38-independent and p38-dependent mechanisms here. Future comparative studies using SB220025 and SB202190 will be useful in identifying the molecular target(s) of SB202190 relevant to cell-type specific vacuole formation.

*BRAF, KRAS* and *EGFR* mutated tumors have been shown to upregulate autophagy for their survival and are amenable to therapeutic autophagy inhibition (26). Interestingly, a recent study could show that non-degradable autophagic vacuoles facilitate apical elimination of RAS-V12 transformed cells, from an epithelial layer of surrounding normal cells (55). This inhibitory role of autophagy in tumor initiation is consistent with increased tumor initiation upon genetic ablation of autophagy in murine models of lung and pancreatic cancer (56, 57). Previous efforts to associate SB202190-sensitivity to oncogenic driver mutations and genotypic alterations in cancer was rather unsuccessful, despite promising effects of SB202190 in suppressing tumor progression in mouse models *in vivo* (24, 25). Expanding the cell-type specificity profile of SB202190 to over 30 cell lines have given us important insights into potential mechanisms of this unique phenotype. Moreover, the differential gene expression analyses have revealed interesting candidates relevant to lysosomal, endosomal and autophagy pathways, which may be used to understand the susceptibility of cancer cell lines to compounds interfering with endosomal/autophagosome maturation. Based on data from The Human Protein Atlas (https://www.proteinatlas.org)(58), the top differentially expressed genes show subcellular localisation corresponding to vesicular compartments (AEBP1, RELN, LAMP5, DLK1, NLRP2, MYOM3) or the cytoplasmic membrane (ANPEP, IGFL2) The knockdown of NLRP2, a gene preferentially expressed in vacuole-forming cells was shown to induce vacuole formation and cell death in HUVECs (Human Umbilical Vein Endothelial Cells) (59). Interestingly, UBE2QL1, a ubiquitin conjugating enzyme less expressed in SB202190-sensitive cell lines was shown to be important to maintain lysosomal integrity and facilitate the autophagic clearance of damaged lysosomes termed lysophagy (60). Monitoring the sensitivity of various cancer cell lines to lysophagy inducers and correlating the responses to the cell-type specificity of vacuolation will be an interesting avenue for future research. Another target, relevant to endosomal transport and downregulated in vacuole forming cells is the lysosome-associated membrane protein-5 gene (LAMP5)(Figure 5C). Originally discovered as brain-associated LAMP-like molecule (BAD-LAMP), LAMP5 has now been shown to play role in haematological and epithelial malignancies (61, 62). Interestingly, LAMP5 was shown to suppress lysosomal degradation of oncogenic fusion proteins promoting mixed-lineage leukemia (61). This autophagy/lysosomal activity suppression by endosomal LAMP5 may be functionally relevant in preventing accumulation of micron-scale vacuoles in non-vacuole forming cells. Lysosomotropic inhibitors of autophagy and lysosomal degradation like chloroquine has provided promising results against cancer (26). The development of more potent and selective lysosomal inhibitors and combined targeting of other kinase signaling pathways is the way forward (63), which requires better understanding of the crosstalks involved.

The vacuoles induced by SB-compounds were originally considered to be autolysosomes and it was assumed that high-basal rates of autophagy may be a cell-type specificity defining factor (25). While autophagy flux analyses in HT29 cells revealed a defective autophagic response associated with LC3-II/p62 accumulation, YFP-LC3 puncta were seen proximal to the micron-scale vacuoles, but did not completely colocalize with them (7). We had previously shown that SB202190 may alter autophagy flux also in non-vacuole forming cells (39). Here we show that while autophagy proficient cells like HT29, HCT116 and A549 show significant alterations in autophagy flux, the vacuole formation is independent of ATG5 and LC3 lipid conjugation as observed in DU-145 and ATG5-deficient A549 cells. This argues against an obligate role for canonical autophagy in SB-induced vacuolation, while autophagy pathway may contribute to the enlarged vesicles in autophagy proficient cells. RAB9 GTPase has been associated with a non-canonical autophagy pathway in the absence of ATG5, but confocal analyses did not reveal consistent colocalization of RAB9 with SB202190-induced vacuoles (Supplementary Figure S1C). In contrast, these compartments are positive for RAB7 GTPase, a late endosomal marker. RAB7-regulated endolysosomal transport has been implicated in tumor progression and RAB7 is an early-Induced lineage and oncogene-specific driver and a prognostic marker in melanoma (64, 65). Confirming potential endosomal origin of these compartments, a recent study had observed RAB7-positive vacuoles in SB203580/SB202190-treated melanoma cells (8). They also indicated similarities between SB202190-induced vacuoles and the vacuolar compartments induced upon the inhibition of the lipid kinase PIKfyve. More recently, another study has suggested PIKfyve as a potential target of SB202190, relevant for vacuolation, based on the similarities in the phenotypes induced by inhibitors of PIKfyve and SB202190 (66). Interestingly, the small molecule inhibitors identified in our screen failed to abrogate vacuolation induced by PIKfyve inhibitors (Supplementary Figure S10). This is supportive of the hypothesis that PIKfyve inhibitors and SB202190 inhibit similar vesicular transport processes, rather than the same target kinase. Consistent with this, a previous study has shown synergistic effect of p38 and PIKfyve inhibitors against cancer cell proliferation *in vitro* and *in vivo* (67). In our hands, the ATR inhibitor VE-821 seems to accurately phenocopy the vacuolation response and cell-type specificity shown by SB202190. Unfortunately, the autophagy-interfering activity of VE-821 and related inhibitors were also shown to be independent of ATR (48). Recently, a CDK4/6 inhibitor Abemaciclib was shown to induce non-canonical cell death associated with LAMP1 positive vacuoles derived from swollen lysosomes (68). Interestingly, like SB202190, despite interference with autophagy pathway, Abemaciclib induced vacuolation does not depend on canonical autophagy. Further studies using VE-821, PIKfyve inhibitors, SB202190 and SB220025 could reveal novel mechanisms and targets responsible for this interesting phenotype relevant to anti-cancer therapy. Methuosis, another micropinocytosis mediated vacuolation process also shows RAB7+ vacuoles and crosstalk with autophagy/lysosomal transport pathways (23, 69). Micropinocytosis inhibitors (70) like dynasore and dyngo-4a does not inhibit SB202190-induced vacuolation in A549 cells (Figure 4A & Supplementary Table S1). However, crosstalk with micropinocytosis may not be ruled out in autophagy-deficient cells. Silmitasertib (CX-4945), a CK2 inhibitor was shown to suppress colon cancer cell growth much like SB202190, but via methuosis rather than autophagy (71). Off note, despite multiple off-targets, several clinical trials have established Silmitasertib as a promising agent against solid tumors, alone and in combination with other chemotherapeutics (72, 73). Multi-target kinase inhibitors like SB202190, Sorafenib, VE-821 and Silmitasertib may be the right tools for anti-cancer intervention rather than specific small molecules. While autophagy is a double-edged sword with respect to anti-cancer interventions, identifying the right nodes of signaling at the interface of lysosomal degradation, endo-lysosomal transport, autophagy and apoptosis will be key to developing effective anti-cancer strategies.

## Supporting information

Supplementary Files-Uncropped blots

## Acknowledgments

MBM thanks Indian Council of Medical Research (ICMR) for support (IRIS #2020-3350). VA was supported by ICMR-Research Associateship (#45/49/2020-Nano/ BMS). SJ, HS and JI thank Department of Education, Government of India for fellowship support and KS acknowledges PhD fellowship support from CSIR, India. SD thanks Department of Biotechnology (DBT), India for Ramalingaswami Re-entry Fellowship (BT/RLF/Re-entry/10/2019) and DBT-India Alliance/Wellcome Trust for Intermediate Fellowship support. The authors would like to thank Kusuma School of Biological Sciences and the Core Research Facility (CRF), IIT Delhi for infrastructural support and confocal microscopy. We would like to thank Drs. Vivekanandan Perumal (IIT Delhi), Ritu Kulshreshtha (IIT Delhi), Archana Singh (CSIR-IGIB), Shantanu Chowdhury (CSIR-IGIB) and Munia Ganguli (CSIR-IGIB) for the gift of cell lines. The authors thank Dr. Matthias Gaestel (MHH, Hannover) for critical reading of the manuscript, Dr. Dirk Heckl (MHH, Hannover) for CRISPR All in one vector and Dr. Amit Tuli (CSIR-IMTECH, Chandigarh) for the gift of GFP-RAB7 plasmids.

## Author contributions

**Susan Jose:** Conceptualization, Methodology, Investigation, Formal analysis **Himanshi Sharma:** Conceptualization, Methodology, Investigation, Visualization, Writing - Review & Editing **Janki Insan:** Conceptualization, Methodology, Formal analysis, Software, Visualization, Writing - Review & Editing **Khushboo Sharma:** Methodology, Investigation, Visualization **Varun Arora:** Methodology, Resources **Sonam Dhamija:** Conceptualization, Methodology, Investigation, Formal analysis, Writing - Review & Editing **Nabil Eid:** Methodology, Resources, Writing - Review & Editing **Manoj B. Menon:** Funding acquisition, Supervision, Conceptualization, Visualization, Writing - Original Draft

## Data Availability

All datasets generated as part of this study are presented here as part of the display items and supplementary files. No new code was generated for the analysis, however the output generated from DESeq2 along with the R scripts can be found at https://github.com/jankinsan/SB202190-dge

