## Supplementary Figures S1-S10 for "Kinase inhibitor-induced cell-type specific vacuole formation in the absence of canonical ATG5-dependent autophagy"

### Supplementary Figure S1.

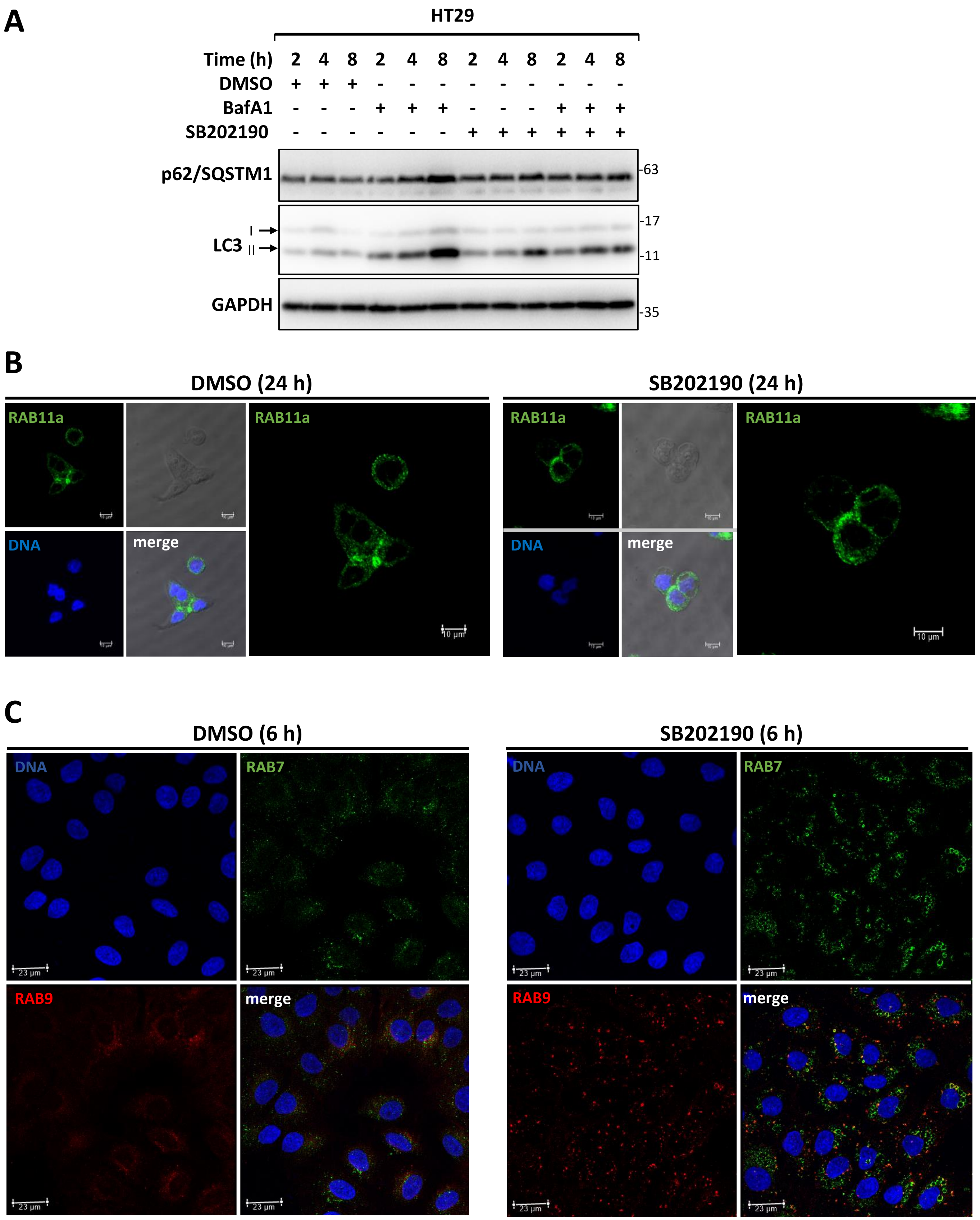

### Supplementary Figure S2.

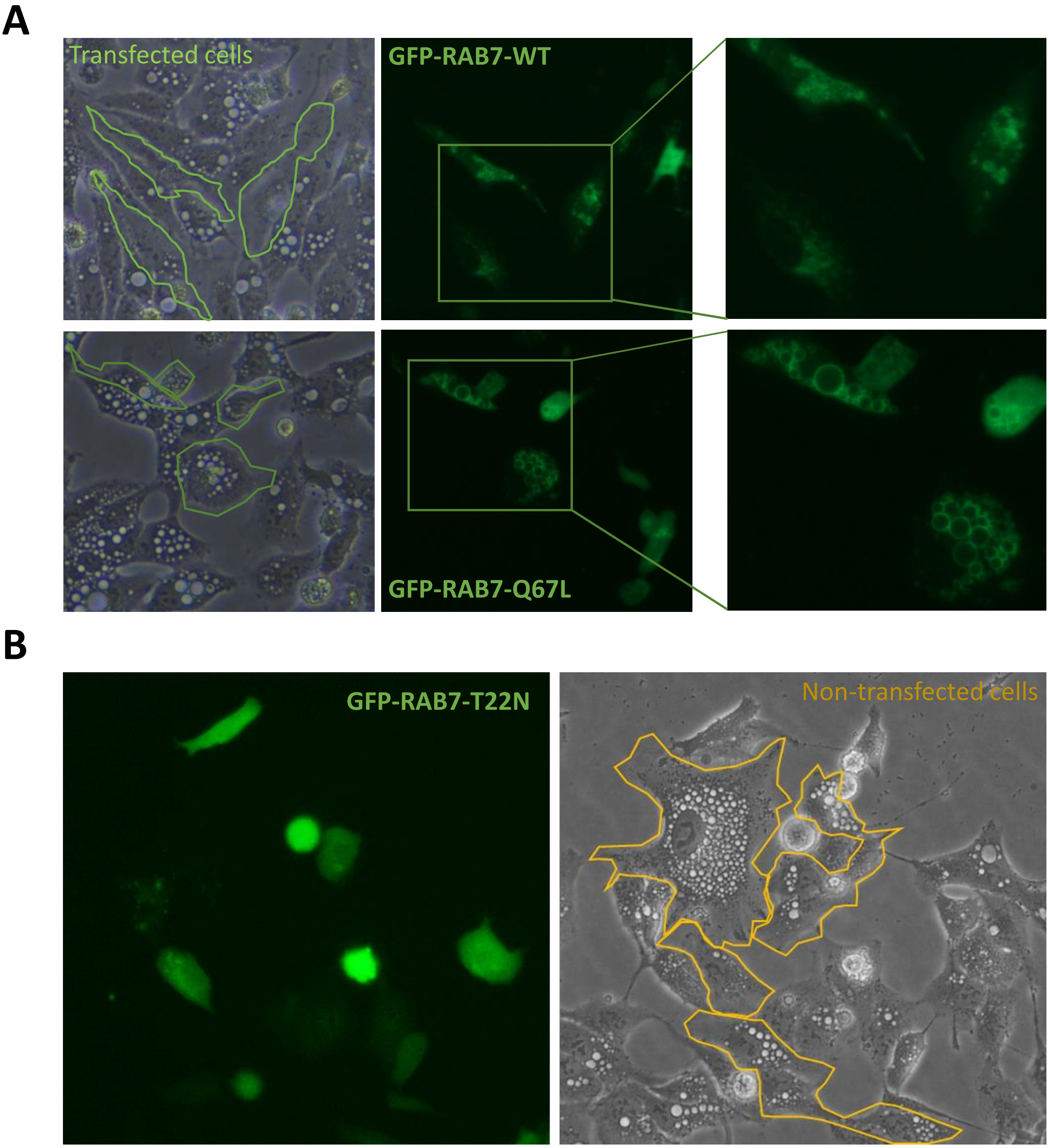

### Supplementary Figure S3.

## A

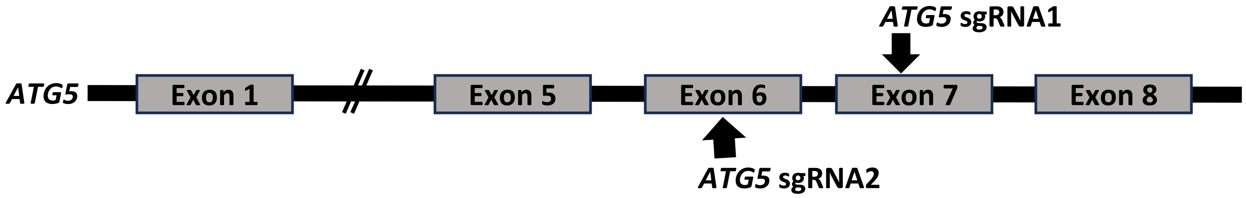


## B

| | | |
|---|---|---|
| *ATG5* sgRNA oligo 1 | Fwd | 5’- CACCGGATGGACAGTTGCACACACT- 3’ |
| | Rev | 5’- AAACAGTGTGTGCAACTGTCCATCC- 3’ |
| Screen Primers (Exon 7) | Fwd | 5’- GAAGGGTCCCAGCCAATTATC - 3’ |
| | Rev | 5’-ATTTCGCCAGTCGAGTCATC - 3’ |

| | | |
|---|---|---|
| *ATG5* sgRNA oligo 2 | Fwd | 5’- CACCGTTCCATGAGTTTCCGATTGA- 3’ |
| | Rev | 5’- AAACTCAATCGGAAACTCATGGAAC- 3’ |
| Screen Primers (Exon 6) | Fwd | 5’- GGCTACTTGAAGCTGTACCTTTG -3’ |
| | Rev | 5’- GAGACAAGAAGGTAGAGCCTCG - 3’ |

## C

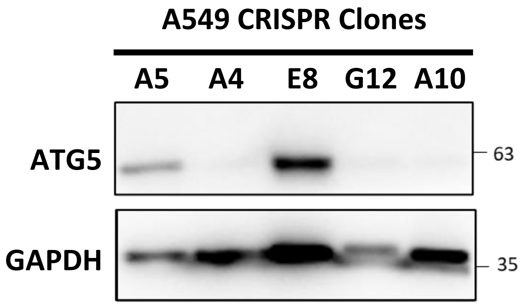

#### Supplementary Figure S4.

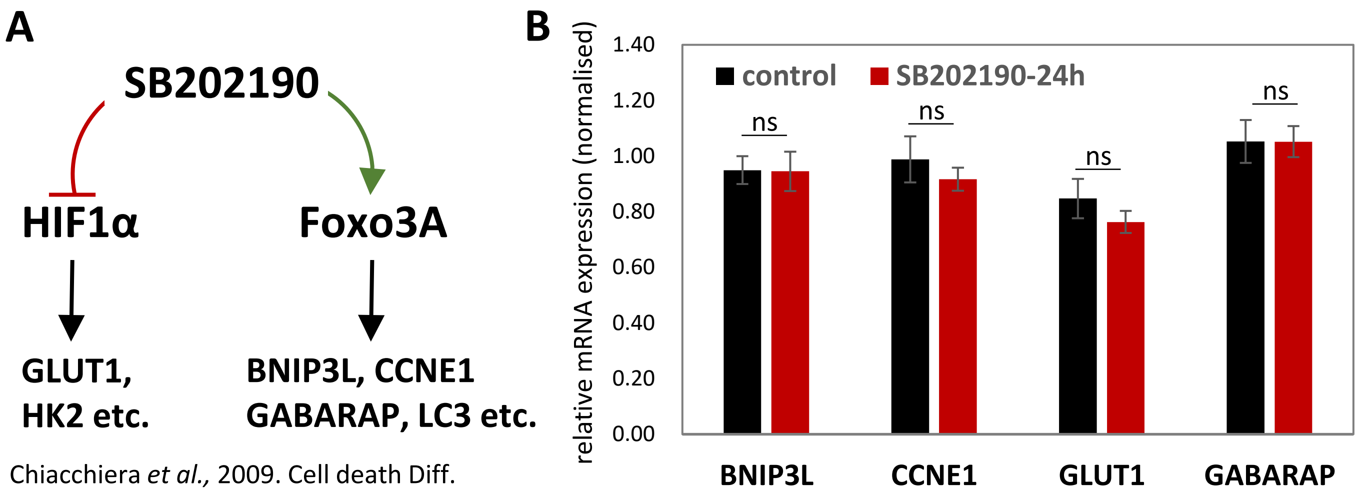

**Supplementary Figure S5.**

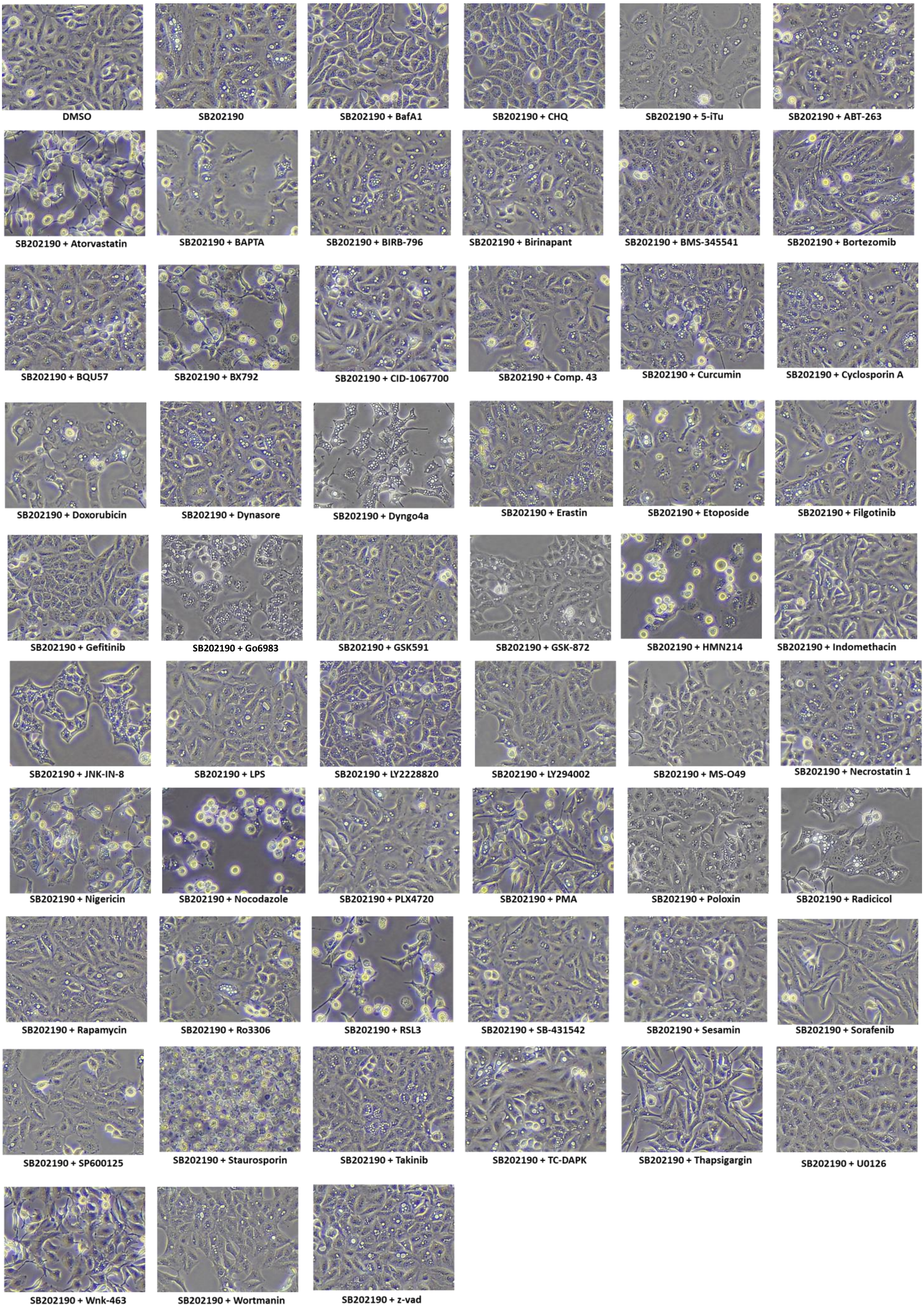

#### Supplementary Figure S6.

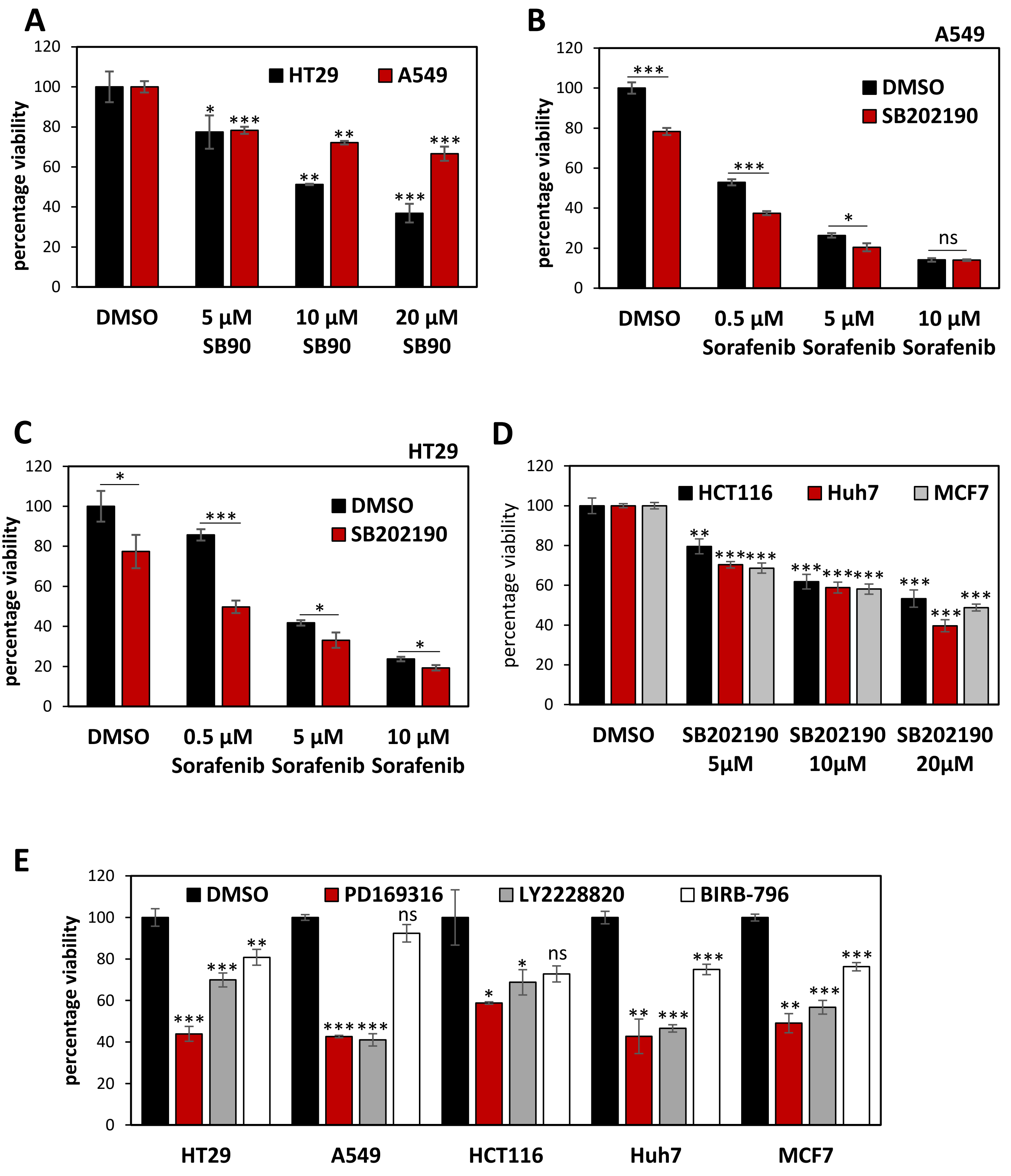

### Supplementary Figure S7.

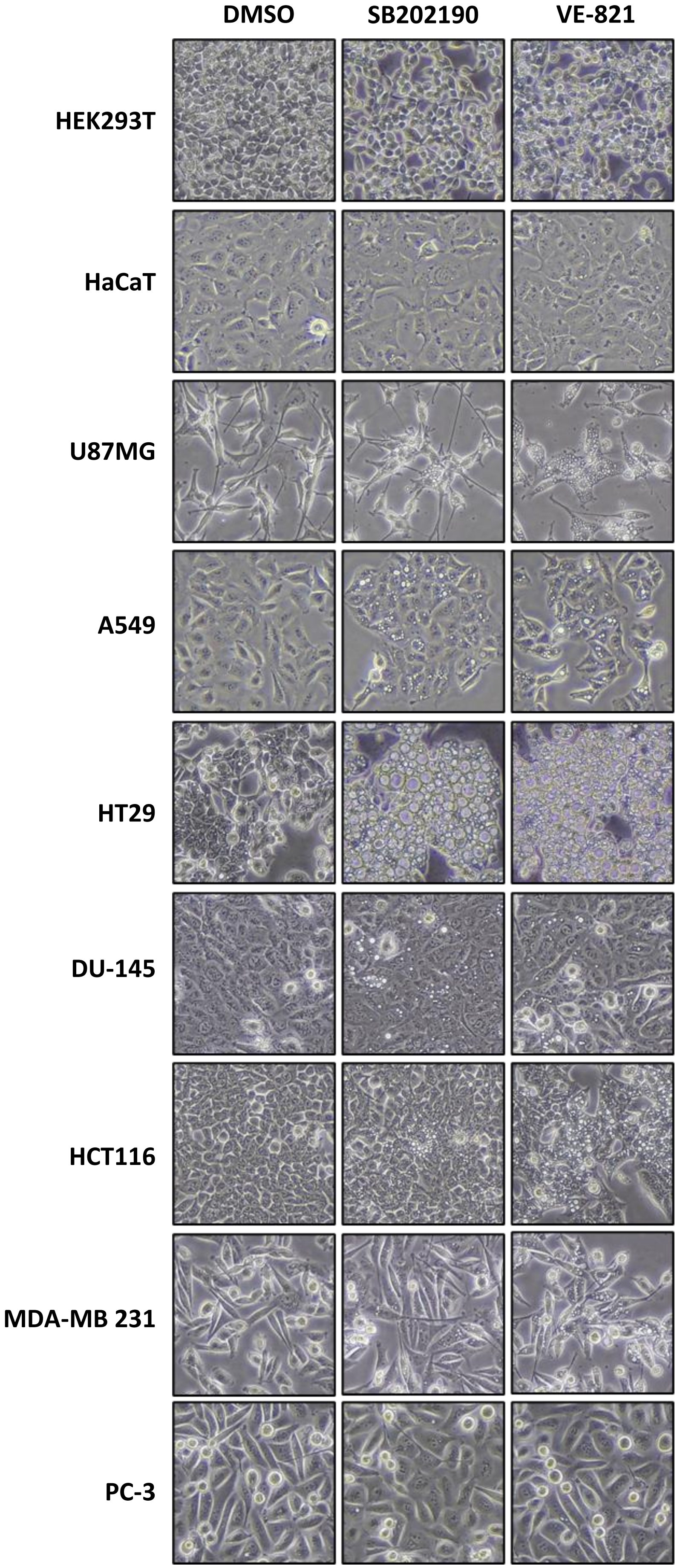

### Supplementary Figure S8.

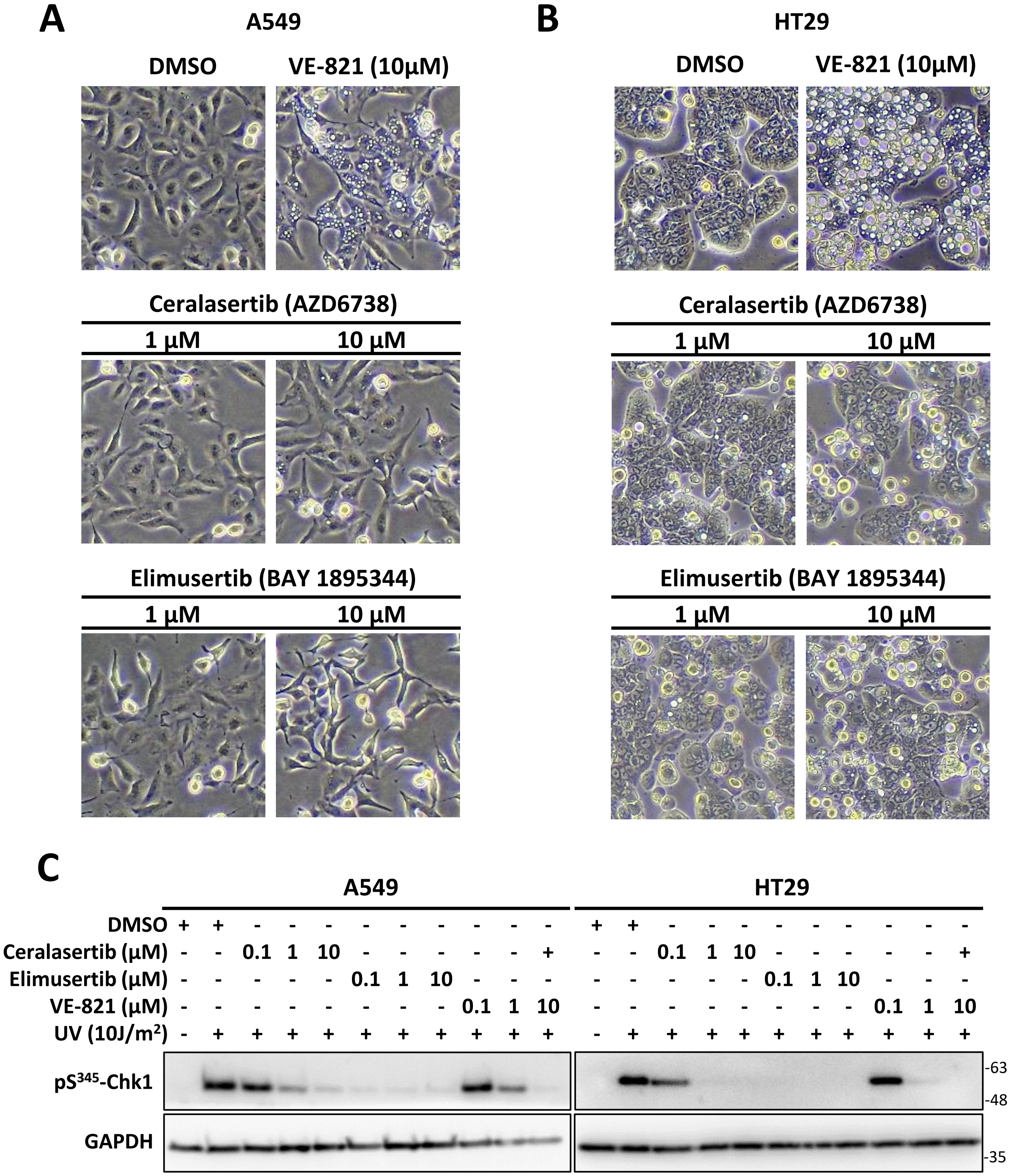

### Supplementary Figure S9.

**A**

**SB202190 Targets**
(Kinomescan - Karaman et al., 2008)

| Gene | Kinase Target | Kd, SB-202190 (nM) |
|---|---|---|
| MAPK14 | p38-alpha | 9.8 |
| NLK | NLK | 28 |
| MAPK11 | p38-beta | 32 |
| MAPK10 | JNK3 | 42 |
| GAK | GAK | 53 |
| CSNK1D | CSNK1D | 59 |
| RIPK2 | RIPK2 | 150 |
| CSNK1E | CSNK1E | 170 |
| RPS6KA1 | RPS6KA1(Kin.Dom.2 - CTD) | 170 |
| EGFR | EGFR(E746-A750del) | 210 |
| MAPK9 | JNK2 | 210 |
| RPS6KA6 | RPS6KA6(Kin.Dom.2 - CTD) | 270 |
| CIT | CIT | 510 |
| PRKACB | PKAC-beta | 530 |
| BRAF | BRAF(V600E) | 620 |
| CDC42BPG | DMPK2 | 640 |
| STK36 | STK36 | 790 |
| EGFR | EGFR(G719C) | 910 |
| ACVR1B | ACVR1B | 950 |
| DDR1 | DDR1 | 1100 |
| EGFR | EGFR(L747-T751del,Sins) | 1100 |
| EGFR | EGFR(L747-S752del, P753S) | 1400 |
| DDR2 | DDR2 | 1600 |
| TNIK | TNIK | 1600 |
| EGFR | EGFR(S752-I759del) | 1700 |
| LATS2 | LATS2 | 1700 |
| PRKACA | PKAC-alpha (PKA) | 1700 |
| CSNK1A1L | CSNK1A1L | 1900 |
| EGFR | EGFR(G719S) | 1900 |
| EGFR | EGFR(L858R) | 1900 |
| RAF1 | RAF1 | 1900 |
| EGFR | EGFR(L747-E749del, A750P) | 2000 |
| EPHA6 | EPHA6 | 2200 |
| EGFR | EGFR(L861Q) | 2300 |
| MAPK8 | JNK1 | 2400 |
| EGFR | EGFR | 2600 |
| GSK3B | GSK3B | 2700 |
| FRK | FRK | 3100 |
| SLK | SLK | 3200 |
| MAPK12 | p38-gamma | 3300 |
| CDC42BPB | MRCKB | 4200 |
| TTK | TTK | 4500 |
| ACVR2B | ACVR2B | 4800 |
| BRAF | BRAF | 4800 |
| ADCK4 | ADCK4 | 4900 |
| ERBB4 | ERBB4 | 4900 |
| STK32B | YANK2 | 4900 |

**VE-821 Targets**
(Millipore KinaseProfiler - Reaper et al., 2011)

| Gene | Kinase Target | % Activity with VE-821 (2μM) |
|---|---|---|
| RAF1 | c-RAF(h) | 11 |
| ROCK2 | ROCK-II(h) | 12 |
| FGFR3 | FGFR3(h) | 14 |
| STK3 | MST2(h) | 15 |
| EPHB4 | EphB4(h) | 17 |
| CAMK2G | CaMKII(r) | 18 |
| MAP2K7 | MKK7beta(h) | 21 |
| AXL | Axl(h) | 22 |
| FYN | Fyn(h) | 22 |
| PRKAA1 | AMPK(r) | 24 |
| CDK1 | CDK1/cyclinB(h) | 25 |
| CSNK1A1 | CK1(y) | 27 |
| NTRK1 | TrkA(h) | 29 |
| RET | Ret(h) | 32 |
| LYN | Lyn(h) | 36 |
| LCK | Lck(h) | 37 |
| CDK5 | CDK5/p35(h) | 43 |
| RIPK2 | RIPK2(h) | 64 |
| VEGFR1 | Flt1(h) | 70 |
| RPS6KA2 | Rsk3(h) | 82 |

**Common Targets**

Casein Kinases

RIPK2

S6 Kinases

RAFs

**B**

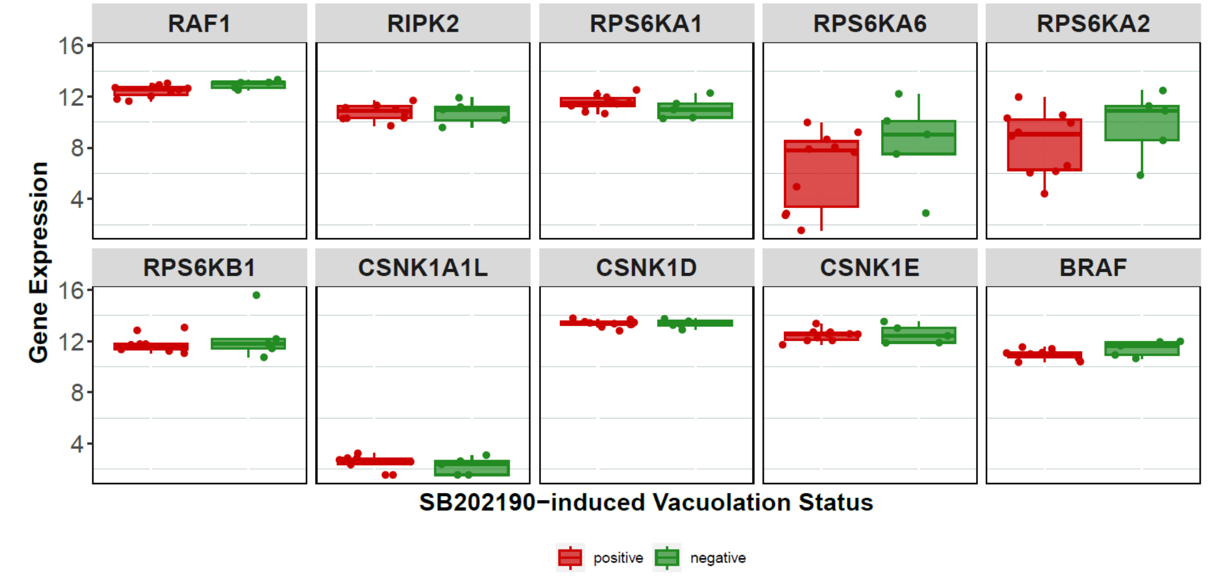

### Supplementary Figure S10.

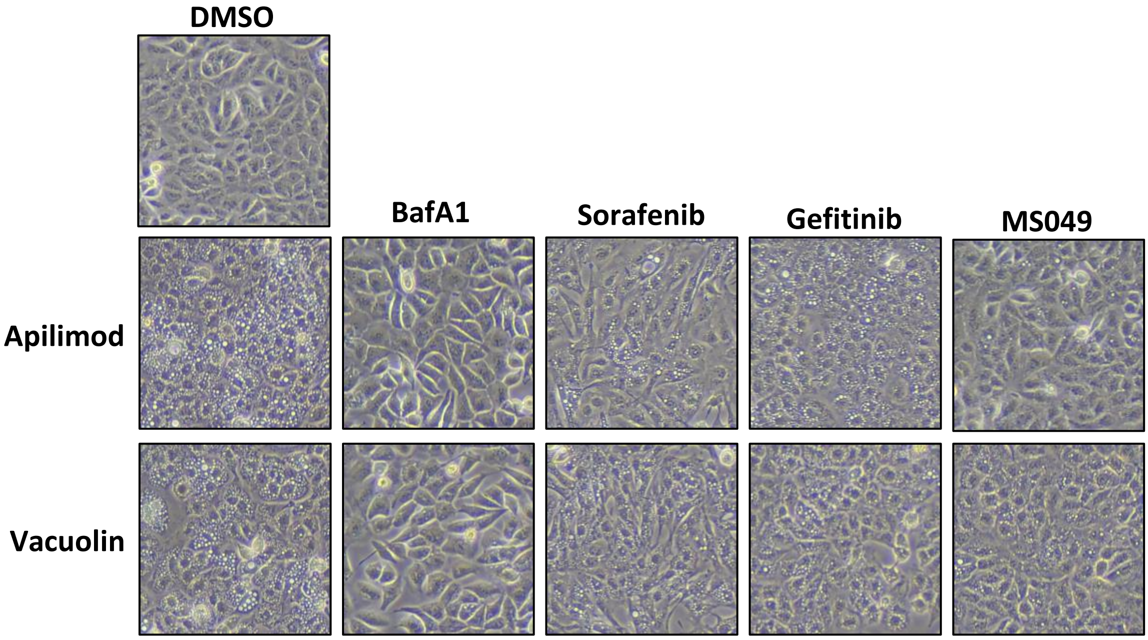
