## Supplementary Information Legends for "Kinase inhibitor-induced cell-type specific vacuole formation in the absence of canonical ATG5-dependent autophagy"

1. **Supplementary Figures S1-S7 (single .pdf file)**

**Supplementary Figure S1. Characterization of SB202190-induced vacuolation. A.** Autophagy flux in SB202190-treated HT29 cells were monitored as in Figure 1D. **B.** HT29 cells treated with SB202190 or DMSO (solvent control) for 24 h and analyzed by confocal immunofluorescence analyses using RAB11a antibodies. DAPI stained nuclei and bright-field images are shown as control. **C.** A549 cells were treated with SB202190 (20µM) or solvent control for 6 h and were analyzed by confocal immunofluorescence analyses using RAB7 (green) and RAB9 (red) antibodies. DAPI was used to stain nuclei.

**Supplementary Figure S2. GTP-bound RAB7 is required for SB202190-induced vacuolation.** A549 cells were transfected with GFP-RAB7-WT (A), GFP-RAB7-Q67L (A) or GFP-RAB7-T22N (B) as indicated and treated with SB202190 for 18 h. **A.** Left panel shows phase contrast images of transfected and treated cells with transfected cells indicated by outline. The right panel shows GFP-RAB7-WT labeled smaller vesicles and GFP-RAB7-Q67L (constitutive GTP-bound mutant) labeled further enlarged vacuoles. **B.** The left panel shows fluorescent images of GFP-RAB7-T22N (non GTP-binding mutant) transfected cells. There is no specific vesicular localization upon SB202190-treatment. On the right panel, phase contrast images show micron-scale vacuoles. Non-transfected cells are indicated, which are predominantly vacuolated, while transfected cells seem to show suppression of SB202190-induced vacuole formation.

**Supplementary Figure S3. CRISPR/Cas9 editing to generate ATG5-deficient A549 cells. A.** Genomic organization of human *ATG5* gene on chromosome 6 with the guide RNA positions indicated. **B.** Sequences of DNA oligos corresponding to the sgRNAs used in the study are shown together with the PCR primers used for amplification and sequencing. **C.** Immunoblot of the single-cell clones showing ATG5-deficient clones A4, A10 and G12. A4 (sgRNA#1) and A10 (sgRNA#2) were used for further analyses.

**Supplementary Figure S4. Effect of SB20190 on gene expression in A549 cells.** A. Gene expression reprogramming induced by SB202190 in colon cancer cells as established by previous studies. B. Realtime qPCR analyses of *BNIP3L, CCNE1, GABARAP* and *GLUT1* expression in control (DMSO) and SB202190-treated (24 h) A549 cells (n=3, ns= not statistically significant with p values >0.05).

**Supplementary Figure S5. Screen for modulators of SB202190-induced vacuolation.** Representative images of A549 cells treated with SB202190 alone or in combination with a panel or inhibitors as indicated for 24 h.

**Supplementary Figure S6. Effect of SB202190 and sorafenib on cell viability. A.** The dose response of SB202190 on cell viability in HT29 and A549 cells treated for 72 h. Statistical significance indicated compared to DMSO control. **B & C.** A549 (B) and HT29 (C) cells were treated with SB202190 (5 µM) alone or in combination with indicated concentrations of sorafenib for 72 h and cell viability was measured by MTT. **D.** The dose response of SB202190 on cell viability in HCT116, Huh7 and MCF7 cells treated for 72 h. Statistical significance indicated compared to DMSO control. **E.** The indicated cell lines were treated with solvent control (DMSO), PD169316 (10 µM), LY2228820 (10 µM) or BIRB796 (1 µM) for 72 h and cell-viability quantified by MTT Assays. Statistical significance indicated compared to DMSO control. n=3, * denotes p<0.05; ** denotes p<0.01 and ***denotes p<0.001 and ns denotes p>0.05.

**Supplementary Figure S7. VE-821-induced cell-type specific vacuole formation.** Various cells lines as indicated were treated with SB202190 (10 µM), VE-821 (10 µM) or solvent control (DMSO) for 24 h and images were acquired.

**Supplementary Figure S8. VE-821 and not other specific ATR-inhibitors induce vacuoles in HT29 and A549 cells. A & B.** A549 (A) and HT29 (B) were treated with solvent control (DMSO) or indicated ATR inhibitors for 24 h and images were acquired to observe vacuolation. **C.** Ceralasertib, VE-821 and Elimusertib are effective ATR inhibitors at the concentrations used, as indicated by the inhibition of UV-induced CHEK1/Chk1 phosphorylation at an ATR target site (Serine-345). GAPDH is shown as loading control.

**Supplementary Figure S9. Expression of common off-targets of VE-821 and SB202190 and sensitivity to vacuolation. A.** Potential direct off-targets of SB202190 and VE-821 were obtained from indicated references and common targets were identified. The common targets included casein kinase isoforms, ribosomal S6-kinase isoforms, RIPK2 and RAF kinases RAF1 and BRAF. **B.** Boxplots display the relative expression levels of these target genes between vacuole-forming and non-vacuolating cancer cell lines as shown in Figure 5. Adjusted p-values are shown and no significant differences were observed.

**Supplementary Figure S10. Vacuoles formed by PIKfyve inhibitors.** A549 cells were treated with PIKfyve inhibitors Apilimod (10 µM) or Vacuolin (10 µM) alone or in combination with the indicated inhibitors for 24 h and images were acquired.

2. **Supplementary Table S1. Inhibitors used in the screen for modulators of SB202190-induced vacuole formation, their working concentrations and effects** (MS Word file).

3. **Supplementary Table S2. Real-time RT-qPCR primer sequences** (MS Word file).

4. **Supplementary Table S3. Genes differentially expressed between vacuole positive and negative cell lines** (MS Excel file)

5. **Supplementary Table S4. Statistical Source data** (MS Excel file)

6. **Supplementary Material-Uncropped Immunoblots** (.pdf document) .
