## Supplementary Table S1 for "Kinase inhibitor-induced cell-type specific vacuole formation in the absence of canonical ATG5-dependent autophagy"

**Supplementary Table 1. Inhibitors used in the screen for modulators of SB202190-induced vacuole formation, their working concentrations and effects**

| **No.** | **Molecule** | **Source** | **Information** | **Concentration** | **Toxicity** | **Morphology change** |
| --- | --- | --- | --- | --- | --- | --- |
| **+** | Bafilomycin A1 | Cayman, Cay11038 | lysosomal acidification inhibitor | 100 nM | + | Yes |
| **+** | Chloroquine | Sigma-Aldrich, C6628 | lysosomal acidification inhibitor | 70 µM |  | Yes |
| 1 | 5-iodo-tubercidin | Medchem Express, HY-15424 | Adenosine kinase inhibitor | 10 µM | + | Yes |
| 2 | ABT-263 | Cayman, Cay11500 | BH3 mimetics | 10 µM |  |  |
| 3 | Atorvastatin | Tokyo Chemical Industry, A2476 | HMGCR inhibitor | 10 µM | ++ | Yes |
| 4 | BAPTA | Santa Cruz,  sc-202488 | Calcium chelator | 10 µM | + |  |
| 5 | Doramapimod (BIRB-796) | Cayman, Cay10460 | p38 (α/β/γ/δ) inhibitor | 1 µM |  |  |
| 6 | Birinapant | Cayman, Cay19699 | SMAC Mimetics | 1 µM |  |  |
| 7 | BMS-345541 | Medchem Express, HY-10518 | IKK1/2 inhibitor | 5 µM |  |  |
| 8 | Bortezomib | LC laboratories,  LC-B-1408 | proteasome inhibitor | 100 nM |  |  |
| 9 | BQU57 | Cayman, Cay16960 | Ral GTPase inhibitor | 10 µM |  |  |
| 10 | BX795 | Tebu-Bio, T1830 | TBK1/IKKε/PDK1 inhibitor | 10 µM | ++ | Yes |
| 11 | CID-1067700 | Medchem Express, HY-13452 | Rab GTPase Inhibitor | 10 µM |  |  |
| 12 | Compound 43 | Cayman, Cay25632 | TAOK1/2inhibitor | 10 µM |  |  |
| 13 | Curcumin | Himedia, RM1449 | Phytochemical | 10 µM |  |  |
| 14 | Cyclosporin-A | Cayman, Cay12088 | calcineurin inhibitor | 2 µM |  |  |
| 15 | Doxorubicin | Geno Bioscience, RC2093 | DNA damage inducer | 10 µM | + | Yes |
| 16 | Dynasore | Sigma-Aldrich, D7693 | Dynamin inhibitor | 10 µM |  |  |
| 17 | Dyngo-4a | Abcam, ab120689 | Dynamin inhibitor | 10 μM | + | Yes |
| 18 | Erastin | Cayman, Cay17754 | Ferroptosis inducer | 10 µM |  |  |
| 19 | Etoposide | Medchem Express, HY-13629 | topoisomerase II inhibitor | 10 µM | + | Yes |
| 20 | Filgotinib | Medchem Express, HY-18300 | JAK1 inhibitor | 10 µM |  | Yes |
| 21 | Gefitinib | Axon, ZD-1839 | EGFR inhibitor | 10 µM |  |  |
| 22 | Go6983 | Sigma-Aldrich, #365251 | PKC inhibitor | 10 μM |  |  |
| 23 | GSK-591 | Medchem Express, HY-100235 | PRMT5 inhibitor | 10 µM |  |  |
| 24 | GSK-872' | MedChem Express HY-101872 | RIPK3 inhibitor | 10 µM |  |  |
| 25 | HMN214 | Medchem Express, HY-12045 | PLK1 inhibitor | 1 µM | +++ |  |
| 26 | Indomethacin | Cayman, Cay70270 | Cyclooxygenase inhibitor | 100 µM |  |  |
| 27 | JNK-IN-8 | Medchem Express, HY-13319 | JNK inhibitor | 10 µM | ++ | Yes |
| 28 | Lipopolysaccharide | Sigma-Aldrich, L6529 | TLR4 agonist | 1 µg/ml |  |  |
| 29 | Ralimetinib dimesylate (LY2228820) | Medchem Express, HY-13241 | p38α/β inhibitor | 10 µM |  |  |
| 30 | LY294002 | Medchem Express, HY-10108 | PI3K inhibitor | 10 µM |  |  |
| 31 | MS-049 | Medchem Express, HY-100360 | PRMT4/6 inhibitor | 10 μM |  |  |
| 32 | Necrostatin 1 | Cayman, Cay11658 | RIPK1 inhibitor | 50 µM |  |  |
| 33 | Nigericin | Cayman, Cay11437 | K^+^/H^+^ ionophore | 10 uM | ++ |  |
| 34 | Nocodazole | Cayman, Cay13857 | Microtubule destabilizer | 50 nM | ++ | Yes |
| 35 | PLX4720 | Cayman, Cay15142 | BRAF inhibitor | 10 µM |  |  |
| 36 | Phorbol-12-myristate-13-acetate | Cayman, Cay10008014 | PKC/MAPK activator | 100 nM |  | Yes |
| 37 | Poloxin | Medchem Express, HY-12134 | PLK1 inhibitor | 10 μM |  |  |
| 38 | Radicicol | Sigma, R2146 | Hsp90 inhibitor | 10 μM | + | Yes |
| 39 | Rapamycin | Tokyo Chemical Industry, R0097 | mTOR Inhibitor | 100 nM |  |  |
| 40 | Ro-3306 | Cayman, 15149 | CDK1 inhibitor | 5 μM |  |  |
| 41 | RSL3 | Medchem Express, HY-100218A | GPX4 inhibitor | 10 µM | ++ |  |
| 42 | SB-431542 | Medchem Express, HY-10431 | ALK5/TGF-β type I Receptor inhibitor | 10 µM |  |  |
| 43 | Sesamin | Medchem Express, HY-N0121 | Delta 5 desaturase inhibitor | 10 µM |  |  |
| 44 | Sorafenib | Medchem Express, HY-10201 | Multi-kinase inhibitor | 10 µM | + | Yes |
| 45 | SP600125 | Medchem Express, HY-12041 | JNK1/2/3 inhibitor | 10 µM |  | Yes |
| 46 | Staurosporine | Medchem Express, HY-15141 | Pan-protein kinase inhibitor | 10 µM | ++++ |  |
| 47 | Takinib | Medchem Express, HY-103490 | TAK1 inhibitor | 10 µM |  |  |
| 48 | TC-DAPK 6 | Medchem Express, HY-15513 | DAP kinase 1/3 inhibitor | 5 µM |  |  |
| 49 | Thapsigargin | Sigma-Aldrich, T9033 | SERCA inhibitor | 1 µM |  | Yes |
| 50 | U0126 | Medchem Express, HY-12031 | MEK1/2 inhibitor | 10 µM |  |  |
| 51 | VE-821 | Cayman, Cay17587-1 | ATM/ATR inhibitor | 10 µM |  |  |
| 52 | Wnk463 | Medchem Express, HY-100626 | WNK inhibitor | 5 µM |  |  |
| 53 | Wortmannin | Sigma-Aldrich, #681675 | PI3K inhibitor | 1 µM |  |  |
| 54 | z-VAD(oMe)-fmk | Medchem Express, HY-16658 | pan-caspase inhibitor | 25 µM |  |  |
