## Supplementary Table S2 for "Kinase inhibitor-induced cell-type specific vacuole formation in the absence of canonical ATG5-dependent autophagy"

| **Gene** | **Primers** | **Sequences (5’-3’)** |
| --- | --- | --- |
| CCNE1 (Cyclin E1) | hCCNE1_Fwd | GGC CAA AAT CGA CAG GAC |
|  | hCCNE1_Rev | GGG TCTG CAC AGA CTG CAT |
| BNIP3L (BCL2 Interacting Protein 3 like) | hBNIP3L_Fwd | GCA CAA CAT GAA TCA GGA CAG |
|  | hBNIP3L_Rev | CAT CTT CTT GTG GCG AAG G |
| GABARAP (GABA Type A Receptor-Associated Protein) | hGABARAP_Fwd | GCG AGA AGA TCC GAA AGA AA |
|  | hGABARAP_Rev | GAT CAG AAG GCA CCA GGT ATT T |
| GLUT1 (Glucose transporter Type I) | hGLUT1_Fwd | GGT TGT GCC ATA CTC ATG ACC |
|  | hGLUT1_Rev | CAG ATA GGA CAT CCA GGG TAG C |

**Supplementary Table S2. Real-time RT-qPCR primer sequences**
