## Supplementary figures and images for "Kinase inhibitor-induced cell-type specific vacuole formation in the absence of canonical ATG5-dependent autophagy"

### Supplementary Files-Uncropped blots

Figure 1D

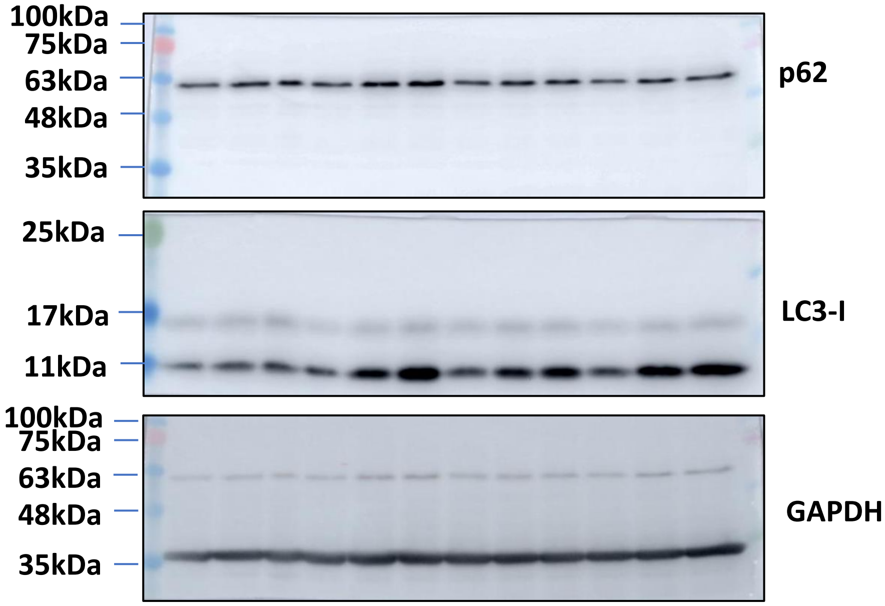


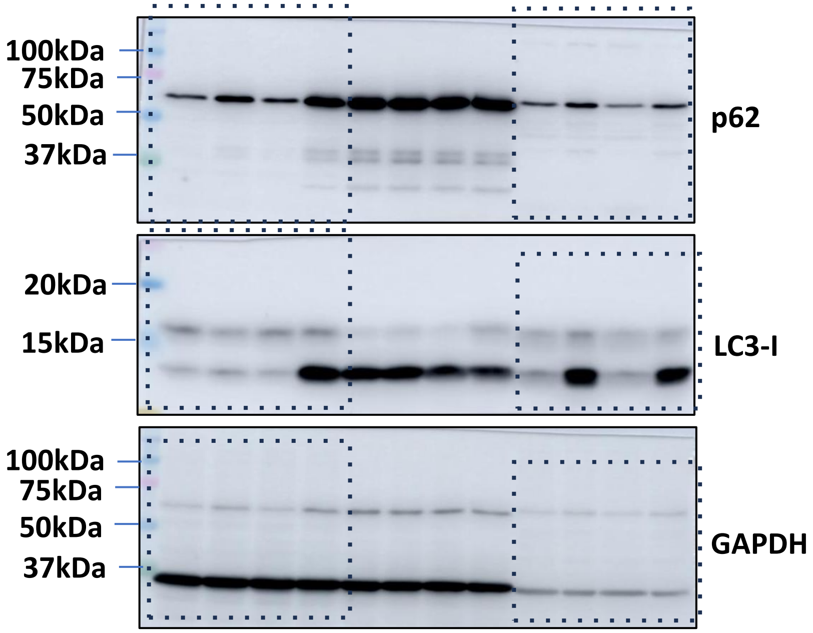


Figure 1E

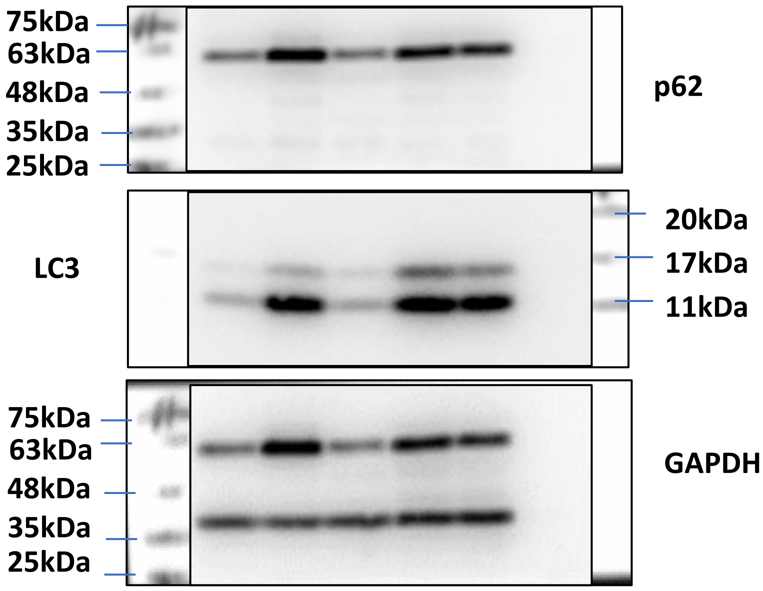

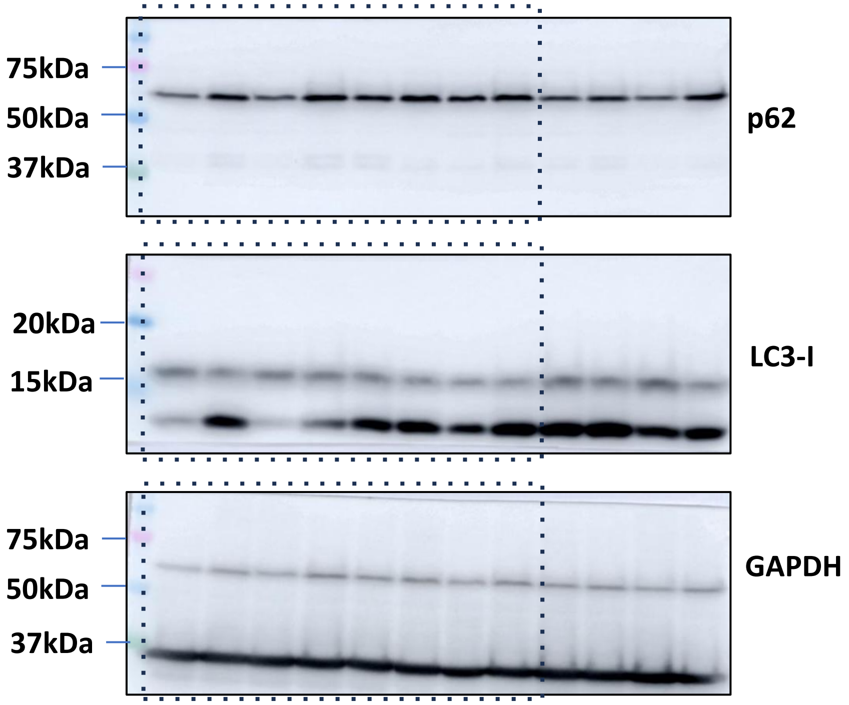

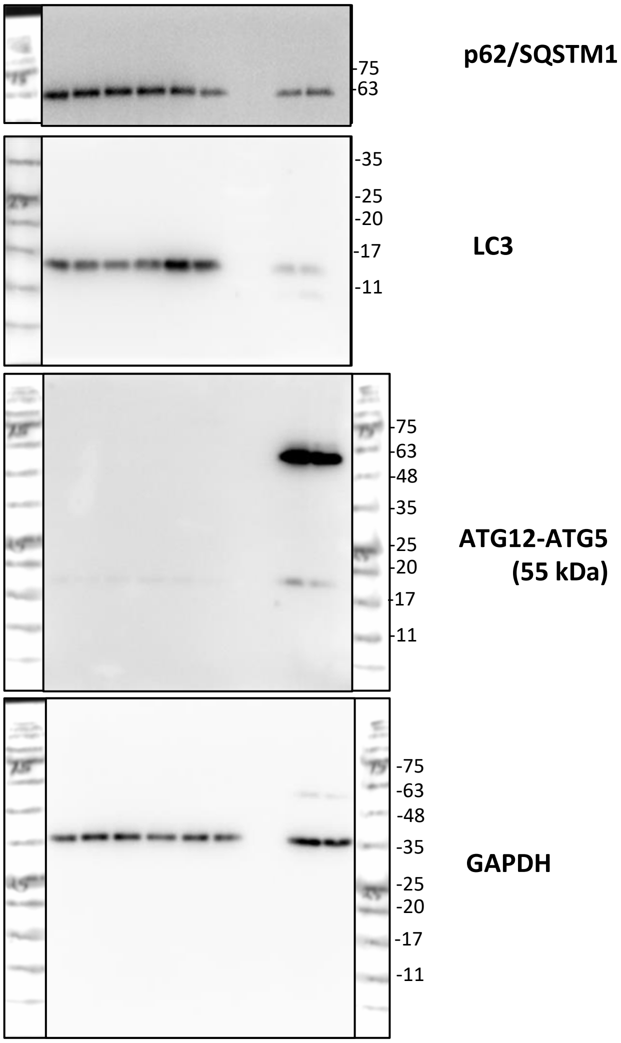

**Figure 3A**

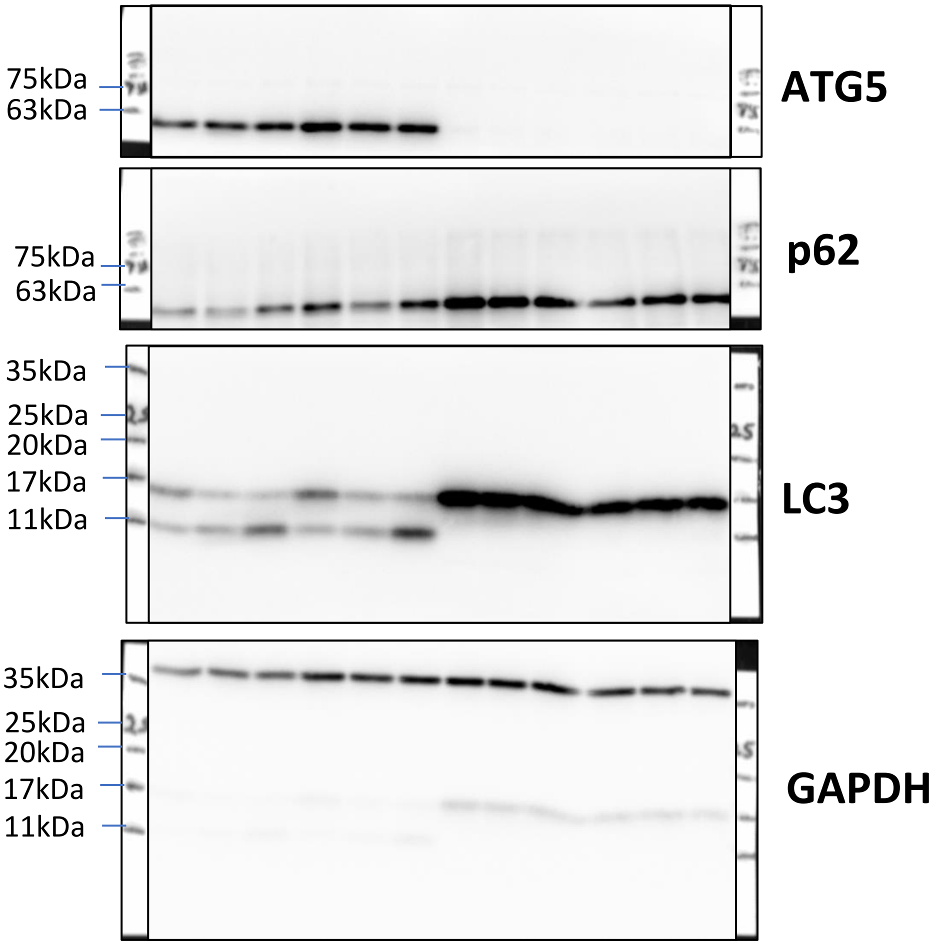


**Figure 3B**

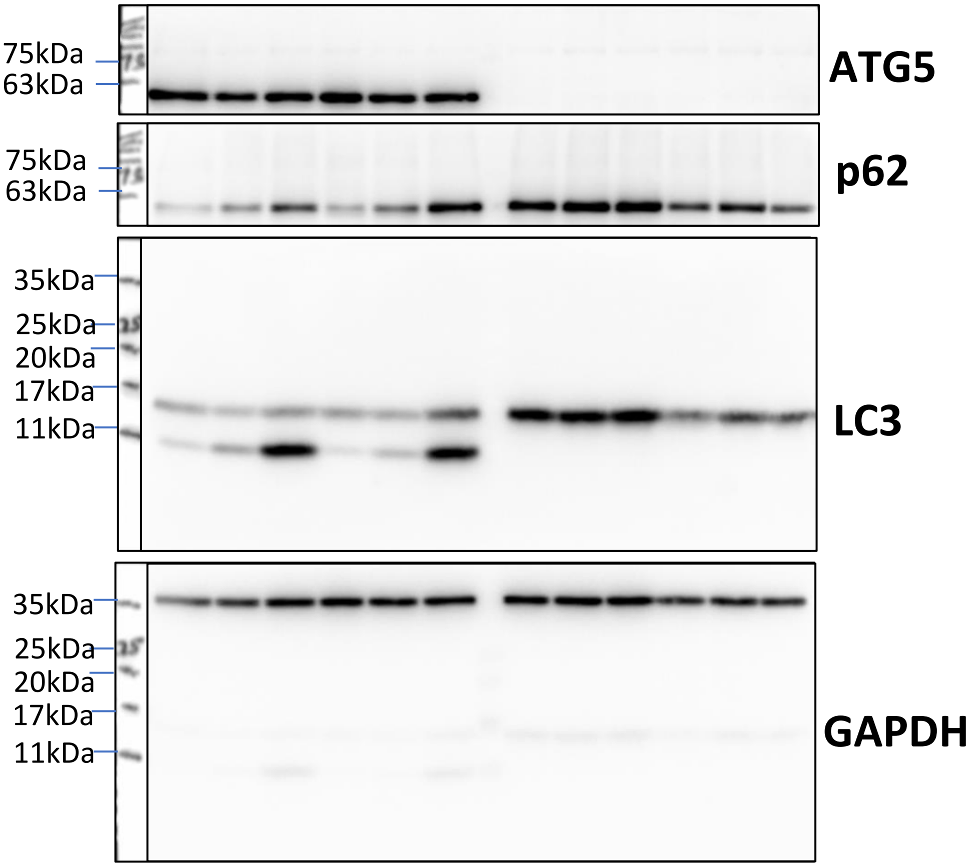

#### Figure S1A

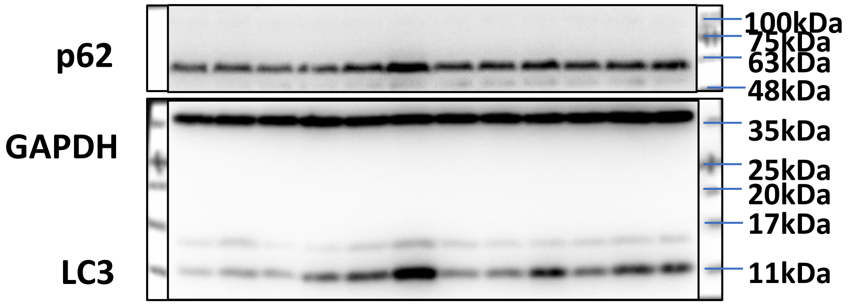


#### Figure S3C

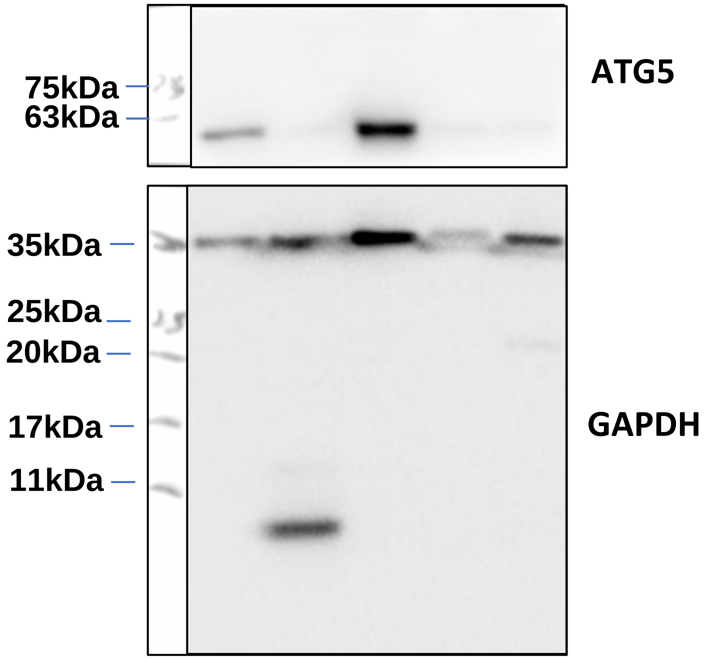


#### Figure S8C

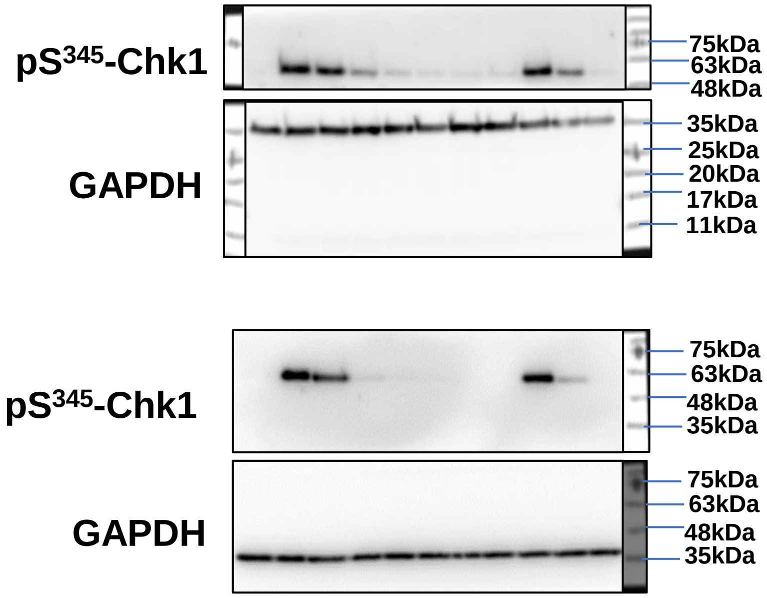
